# DNA hypomethylation activates Cdk4/6 and ATR to cause dormant origin firing and cell cycle arrest that restricts liver outgrowth in zebrafish

**DOI:** 10.1101/2023.06.26.545878

**Authors:** Bhavani P. Madakashira, Elena Magnani, Shashi Ranjan, Kirsten C. Sadler

## Abstract

Coordinating epigenomic inheritance and cell cycle progression is essential for organogenesis. UHRF1 connects these functions by facilitating maintenance of DNA methylation and cell cycle progression. Here, we provide evidence resolving the paradoxical phenotype of *uhrf1* mutant zebrafish embryos that have both activation of pro-proliferative E2F target genes and increased number of hepatocytes in S-phase, but the liver fails to grow. We find that Atr inhibition reduces DNA replication and increases liver size in *uhrf1* mutants, suggesting that *uhrf1* mutant hepatocytes have replication stress leading to Atr-mediated cell cycle inhibition and dormant origin firing. We uncover persistent Cdk4/6 activation as the mechanism driving *uhrf1* mutant hepatocytes into S-phase, activating Atr and restricting hepatic outgrowth. Palbociclib treatment of *uhrf1* mutant embryos prevented aberrant S-phase entry, and the DNA damage response. Palbociclib rescued most cellular and developmental phenotypes in *uhrf1* mutants, except DNA hypomethylation, transposon activation and the interferon response. Pro-proliferative genes were also activated in a Cdk4/6 dependent fashion in the liver of *dnmt1* mutants, suggesting DNA hypomethylation as a mechanism of Cdk4/6 activation. Thus, the developmental defects caused by DNA hypomethylation are attributed to persistent Cdk4/6 activation leading to DNA replication stress, dormant origin firing and cell cycle inhibition, preventing hepatic outgrowth.

## Introduction

Cell cycle progression during organogenesis requires precise coordination of the machinery that replicates the genome and those that pattern the epigenome. The chromatin landscape dictates the timing of regions of the genome get replicated early or late in S-phase and can influence when and where the origins of DNA replication fire ^1^. Thus, the widespread changes to the epigenome that occurs during development when cell identity is established, in aging or in cancer can influence the pattern, timing or progression of DNA replication ^2, 3^. The crosstalk between the pathways that control cell cycle progression and epigenetic patterning during development has not been well studied.

DNA methylation is among the best studied epigenetic marks. The methylome is patterned during early vertebrate development and is maintained during cell division by a process that couples DNA replication to DNA methylation via interaction between proteins at the replication fork. Newly synthesized DNA is unmethylated and the hemimethylated structure of DNA generated at the replication fork is recognized by Ubiquitin-like, containing PHD and RING finger domains, 1 (UHRF1) which, through a base flipping mechanism ^4–7^, allows DNA methyltransferase 1 (DNMT1) to access and methylate the daughter strand ^8–10^. Both proteins are anchored to the DNA replication fork ^1^^1, 12^ through interactions with PCNA ^10^, LIG ^13^ and modified histones ^10, 14, 15^. Thus, the colocalization and coactivity of the methylation and replication factors assures faithful propagation of methylation patterns.

One way to assure that these factors act together is by co-regulating their expression. UHRF1, DNMT1 and all of the genes required for DNA replication are induced by the E2F family of transcription factors ^16–20^. E2F is activated when the retinoblastoma (Rb1) protein becomes phosphorylated by the cyclin-dependent kinases 4/6 (CDK)-Cyclin D complex. Therefore, the downstream consequence of CDK4/6 activation in response to mitogen stimulation assures that the factors required for both DNA replication and DNA methylation are present as cells enter S-phase. This coupling is essential, as in both during development and in cancer cells, inactivation of either UHRF1 or DNMT1 blocks cell proliferation ^21–29^ by a mechanism that is not understood.

Just as these cell cycle regulatory pathways can regulate DNA methylation, the chromatin landscape influences the cell cycle. For instance, repressive epigenetic marks restrict E2F target gene accessibility, and removal of these repressive marks is essential for E2F function ^30^. The p16^INK4a^ gene, which encodes a CDK4/6 inhibitor, is regulated by repressive epigenetic marks which can lead to suppression of p16^INK4a^, causing inappropriate cell cycle entry ^31^. Another study showed that with DNMT1/DNMT3B deficient cells, loss of 3D genome structure randomized replication timing ^32^. These data demonstrate the complexity of the interplay between the ubiquitous cell cycle machinery and epigenetic modifiers to ensure fidelity of the epigenetic patterns as cells proliferate. There is a limited understanding of how these processes are regulated during the rapid cell cycles that take place during vertebrate development.

The developing liver provides an optimal model to study this relationship, as liver outgrowth is mediated by proliferation of differentiated hepatocytes. In zebrafish, hepatocytes are specified and determined during the organogenesis stage of development (between 24-48 hours post fertilization, hpf) ^33, 34^ and the differentiated hepatocytes proliferate while the liver undergoes morphogenesis during hepatic outgrowth. This stage is largely complete by 120 hpf ^35^. There are a few studies that indicate that the mechanisms of cell cycle regulation during hepatic outgrowth utilize the same signaling pathways that control proliferation in most other cell types. For instance, in mice, genetic approaches showed that E2F regulates hepatocyte proliferation during liver development and regeneration ^36, 37^ and hepatocyte specific knockout of Rb1 is sufficient to promote cell cycle entry, increased ploidy and generated large nuclei in post-natal and adult mouse hepatocytes ^38, 39^. It is not known if these mechanisms are conserved in zebrafish or how these and other critical cell cycle regulators control organ growth.

We previously discovered a functional relationship between *uhrf1* and cell cycle progression during hepatic outgrowth in zebrafish. *uhrf1* mutant embryos have multiple developmental abnormalities, including a small liver ^23, 26, 40^. These phenotypes are first evident between 80-96 hpf and are exaggerated at 120 hpf, resulting in lethality by 10 days post fertilization (dpf) ^25^. The cellular phenotypes of *uhrf1* deficient hepatocytes are similar to those of *dnmt1* mutants: mutant larvae have large, misshapen nuclei, near complete loss of nuclear structures ^41^, and widespread cell death ^23, 25^, which is dependent on Tnfa ^42^. Most strikingly, at 120 hpf, when DNA replication is complete in wild-type larvae (WT), nearly half of all mutant hepatocytes are undergoing DNA replication with a gene expression signature typical for proliferating cells^25^ despite the fact that the liver does not grow. As in all other systems studied ^8, 9, 26, 43, 44^, *uhrf1* and *dnmt1* loss in zebrafish results in DNA hypomethylation, which unleashes transposons and an interferon response to these viral mimetics ^42, 45^. It is not known which cellular and molecular defects contribute to the developmental phenotypes of these mutants.

Here, we resolving the paradoxical finding of the cellular phenotype and molecular phenotypes associated with DNA replication and proliferation with the developmental phenotype of hepatic outgrowth failure in *uhrf1* mutants. We identify Cdk4/6 activity as the mechanism giving rise to most of the developmental and cellular defects in *uhrf1* zebrafish mutants, except for DNA hypomethylation, transposon activation and induction of the interferon response. The Cdk4/6 inhibitor, Palbociclib, rescues all embryonic morphological defects in *uhrf1* mutants, including liver size, DNA rereplication, nuclear enlargement and euchromatinization and cell death. Strikingly, embryo survival is improved by Palbociclib. Cdk4/6 signaling is also activated in another model of DNA hypomethylation, in *dnmt1* mutants, suggesting that DNA hypomethylation as the underlying mechanism. Since DNA replication does not occur in *dnmt1* mutants and Dnmt1 depletion in *uhrf1* mutant hepatocytes blocks DNA replication, we concluded that Dnmt1 is essential at the replication fork ^11, 12^. The Ataxia Telangiectasia And Rad3 Related (Atr) pathway is activated in *uhrf1* mutants, signifying replication stress and we show that DNA replication in part requires Atr signaling. These data provide new mechanistic insight into the interaction between key cell cycle regulatory pathways and DNA methylation during development, indicate the importance of the Cdk4/6 pathway in cellular consequences of Uhrf1 depletion. This also suggests that the important chemotherapeutic, Palbociclib, may modulate the cellular response to epigenetic defects.

## Results

### DNA rereplication is a primary cellular response to *uhrf1* loss

We previously reported that *uhrf1* mutant livers have an increased number of cells that incorporate nucleotides during hepatic outgrowth, despite the livers being smaller sized ^25^. To determine whether this was due to DNA replication or DNA repair, we investigated the pattern, timing and outcome of EdU incorporation in hepatocytes. FACS sorted hepatocytes from *uhrf1* and WT larvae at 120 hpf showed that 7% of *uhrf1* mutants had 4N DNA content compared to 2% of controls (Figure 1A), consistent with our previous finding of enlarged hepatocyte nuclei in *uhrf1* mutants ^41^, a phenotype that reflects increased ploidy ^46^. We pulsed embryos at 80, 96 and 120 hpf for 30 minutes with EdU, sorted *uhrf1* mutants based on morphological phenotypes at 96 hpf and 120 hpf, and, since the phenotype is not apparent at 80 hpf, all embryos at this time point were individually genotyped and assessed for the number of cells with EdU incorporation. Nearly all hepatocytes in *uhrf1* mutants and their phenotypically normal sibling control larvae (heretofore referred to as controls) were EdU positive at 80 hpf. At 96, and 120 hours, this was reduced to 25% and 10%, respectively in controls. However, 55% and 43% of *uhrf1* mutant hepatocytes were EdU positive at these same time points (Figure 1B-C).

**Figure 1.** DNA rereplication leads to increased DNA content and heterochromatin loss in *uhrf1* mutant hepatocytes. (A) FACS sorting of 5 dpf *Tg(fabp10a:nls:mcherry)* hepatocytes with control or *uhrf1*^-/-^ at 120 hpf. (B) Representative Z-stack projection confocal images of EdU and Hoechst stained control and *uhrf1*^-/-^ livers. Whole control and *uhrf1*^-/-^ embryos were pulsed with EdU for 30 minutes at 80 hpf, 96 hpf and 120 hpf and processed for EdU fluorescence by Click-it. (C) Quantification of EdU incorporation in B. (D) 80 hpf, 96 hpf and 120 hpf hepatocytes were sorted into early, mid and late S phase based on EdU pattern and quantified. (E) Representative immunofluorescent images of 120 hpf *uhrf1*^-/-^ and control livers with heterochromatin markers H3K9me3, H3K9me2 and Heterochromatin and their quantification. (F) Quantification of EdU incorporation from a 12 minute EdU pulse at 80 hpf and 120 hpf control and *uhrf1*^-/-^ hepatocytes and (G) sorted based on their S phase stage. C indicates control siblings and M indicates *uhrf1*^-/-^. Scale bar = 50 µm, the number of samples and clutches indicated for each condition and represented as median with range. p-value *< 0.05, **< 0.005, *** < 0.0005 by unpaired Students t-test with adjustment for multiple comparisons.

The stereotypical pattern of DNA replication has euchromatic DNA replicated in early S-phase, visualized as homogeneous EdU labeling in the nucleoplasm, followed by replication of DNA in the nuclear lamina during mid-S-phase and heterochromatin replicated last ^47^. If the increase in EdU incorporation in *uhrf1* mutant hepatocytes reflected DNA rereplication, then these patterns should be apparent. Scoring S-phase stage based on the patterns of EdU incorporation showed that at 80 hpf, most hepatocytes in controls and *uhrf1* mutants were in early S-phase, while at 96 hpf and 120 hpf there was a significant increase in the number of *uhrf1^-/-^* hepatocytes in mid and late S-phase, respectively (Figure 1D). These data rule out the possibility that EdU incorporation in *uhrf1* mutant hepatocytes reflects DNA damage repair, as repair mediated nucleotide incorporation does not mirror the stereotypical stages of DNA replication ^48, 49^. Interestingly, we observed that *uhrf1^-/-^* hepatocytes incorporate EdU in a pattern that reflects a pattern of heterochromatin, even though these hepatocytes are completely devoid of the heterochromatin markers H3K9me3, H3K9me2, Hp1α (Figure 1E) and the nuclear lamina marker Lamin A ^41^. We conclude that increased number of EdU positive hepatocytes in *uhrf1* mutants reflects DNA rereplication.

We hypothesized that the EdU incorporation phenotype in *uhrf1* mutant hepatocytes reflected either an acceleration of the fork progression rate or an increase in the number of active replication origins. To test this, we optimized the duration of the EdU pulse to be just below the limit of detection in control hepatocytes at 120 hpf (Figure S1). Following a 12 minute pulse with EdU, no hepatocytes were labeled at either 80 or 120 hpf in control larvae or in *uhrf1^-/-^* larvae at 80 hpf but nearly 50% of *uhrf1* mutant hepatocytes were labeled at 120 hpf (Figures 1F and Figure S1), and nearly all of these EdU positive *uhrf1^-/-^* nuclei were in early S-phase (Figure 1G). This suggests that Uhrf1 deficiency either induces firing of more DNA replication origins or increases replication fork speed.

### CDK4/6 is activated in *uhrf1* mutant livers

To identify the basis of the DNA replication defect, we analyzed bulk RNAseq data from pools of control and *uhrf1* mutant livers collected at 120 hpf ^42^, when the hepatic developmental and cellular phenotypes are fully penetrant. Gene ontology (GO) analysis using REVIGO revealed that downregulated genes were mainly involved in hepatic functions, such as protein modification and lysis, signaling and metabolic process, while the upregulated genes were enriched in pathways involved in cell cycle regulation, DNA replication, anti-viral and immune response and cell death (Figure 2A, Table S1). UPSET analysis of the differentially expressed genes (DEGs) in all of the cell cycle pathways identified by REVIGO demonstrated that many genes were common to multiple genesets, however a cluster of genes involved in DNA replication and cell division were unique (Figure 2B; Table S1). These data confirm and extend our previous findings ^23, 25, 42^ that *uhrf1* mutant livers have aberrant cell cycle regulation.

**Figure 2.**
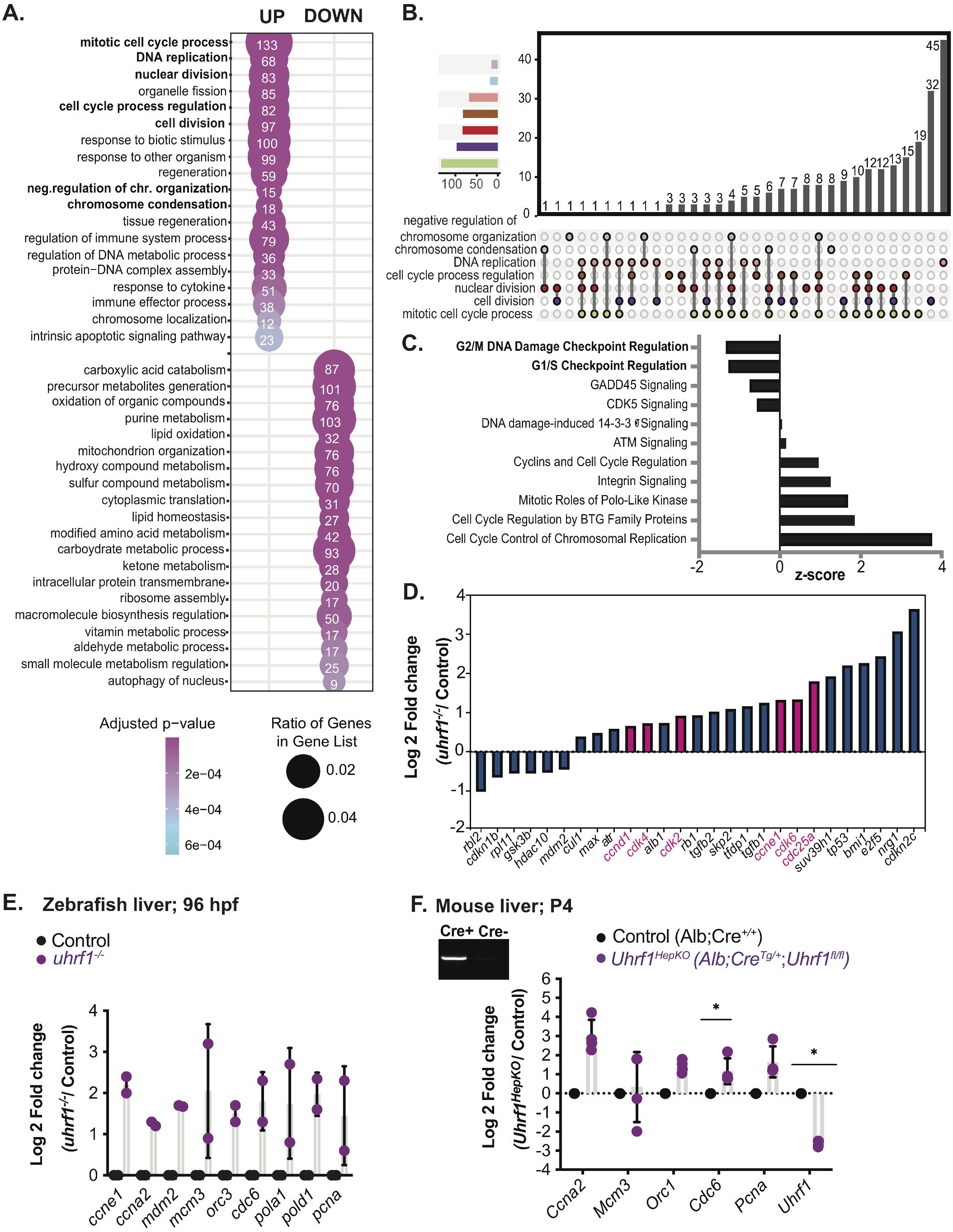
CDK4/6 is activated in *uhrf1* mutant livers. (A) Gene ontology (REVIGO) of the significant upregulated and downregulated genes in 120 hpf *uhrf1*^-/-^ liver RNA-Seq (*p_adj_* < 0.05) for each category. The cell cycle GO terms are highlighted in bold. (B) UPSET plot of all the cell cycle GO category genes from A. (C) Ingenuity Pathway Analysis (IPA) of all the cell cycle genes differentially expressed in 120 hpf *uhrf1*^-/-^ liver RNA-Seq, pathways involved in cell cycle checkpoint regulation in bold, z-score in IPA indicates a predicted activation or inhibition of a pathway. (D) Genes from the G1/S checkpoint pathway generated by IPA plotted with log2fold change, important cell cycle genes highlighted in red. (E) qPCR on key cell cycle genes of 96 hpf *uhrf1*^-/-^ and control livers. (F) qPCR on key cell cycle genes in the livers of P4 neonatal mice deficient for *Uhrf1* in hepatocytes (*Uhrf1^HEPKO^*) and controls, inset shows band for Cre transgene by RT-PCR. The number of clutches is indicated for each condition and represented as median with range. p-value *< 0.05, **< 0.005, *** < 0.0005 by unpaired Students t-test with adjustment for multiple comparisons.

We next used Ingenuity Pathway Analysis (IPA) of the unified set of all DEGs in the GO term genesets related to the cell cycle to identify potential upstream regulators of the cell cycle DEGs in *uhrf1* mutant livers. This uncovered a pattern whereby the G1/S and G2/M checkpoint pathways were downregulated and cell cycle regulation, proliferation and transition through the cell cycle were upregulated (Figure 2C, Table S2). The key upstream regulators of the G1/S transition and S-phase entry and progression, *Cdk4/6-Cyclin D, Cdk2-Cyclin E* and the DNA replication initiation factor, *Cdc25a*, were upregulated in *uhrf1* mutant livers. Additionally, the cell cycle inhibitor *cdkn1b* and the cell cycle regulated gene *rbl2,* which prevents access of E2F1 and is a UHRF1 interacting protein ^50^, were downregulated (Figure 2D). Together, these data suggest that loss of *uhrf1* results in repression of G1/S and G2/M cell cycle checkpoints and activation of Cdk4/6 downstream targets that promote G1/S transition and DNA replication.

These data establish a developmental timeline of the *uhrf1* mutant hepatic phenotypes whereby DNA hypomethylation ^42^, and nuclear size defects are first detected at 80 hpf ^41^, followed by DNA rereplication (Figure 1B-D,F) and cell death at 96 and 120 hpf. Since no cell cycle DEGs were detected at 80 hpf ^25^ and qPCR analysis on RNA from pooled livers dissected at 96 hpf showed that several cell cycle regulatory genes (*ccne1, ccna2)*, replication licensing factors (*mdm2, mcm3, orc3, cdc6)* and processivity factors (*pola1, pold1* and *pcna)* were upregulated at this time point (Figure 2E), we conclude that the DNA replication defect in uhrf1 hepatocytse occurs following loss of DNA methylation. Importantly, we found a similar effect in the liver of neonatal mice at P4 that are deficient for *Uhrf1* in hepatocytes ((*Alb:Cre^Tg/+^;Uhrf1^fl/f^ i.e. Uhrf1^HepKO^*) ^44^. There was robust expression of Cre (Figure 2F, inset) and downregulation of *Uhrf1*, indicating effective Cre-mediated Uhrf1 depletion in these samples. All cell cycle genes analyzed showed increased expression in *Uhrf1^HepKO^* livers (Figure 2F), suggesting that Uhrf1 plays a conserved role in regulating the cell cycle during hepatic outgrowth by promoting S-phase entry and DNA rereplication.

### Cdk4/6 activation is required for the *uhrf1* mutant phenotype

We hypothesized that Cdk4/6 and E2F activation combined with loss of G1/S checkpoints could drive Uhrf1 deficient cells into S-phase. To test this, we used the Cdk4/6 inhibitor Palbociclib (PD in figures), which is an approved chemotherapeutic for breast cancer ^51^ that effectively suppresses DNA replication and cell proliferation in liver cancer cells ^52^. We optimized the dose and timing of Palbociclib exposure to obtain the maximal tolerable concentration and administration timing to achieve no effects on control larvae based on survival, overall embryo phenotype, morphological criteria and liver size (Table S3). The final treatment scheme outlined in Figure 3A involved adding 20 μM Palbociclib in 0.5% DMSO or 0.5% DMSO as a control to 48 hpf embryos generated from an incross of *uhrf1^+/-^* adults which expressed nuclear localized signal targeted cherry transgene in hepatocytes (*Tg(fabp10a: nls-mcherry)*) to mark the livers. The resulting larvae were collected at 120 hpf, imaged to assess morphological features characteristic of *uhrf1* mutants and then all larvae were individually genotyped. In some experiments, larvae at 120 hpf were pulsed for 30 minutes with EdU to assess DNA replication.

**Figure 3.**
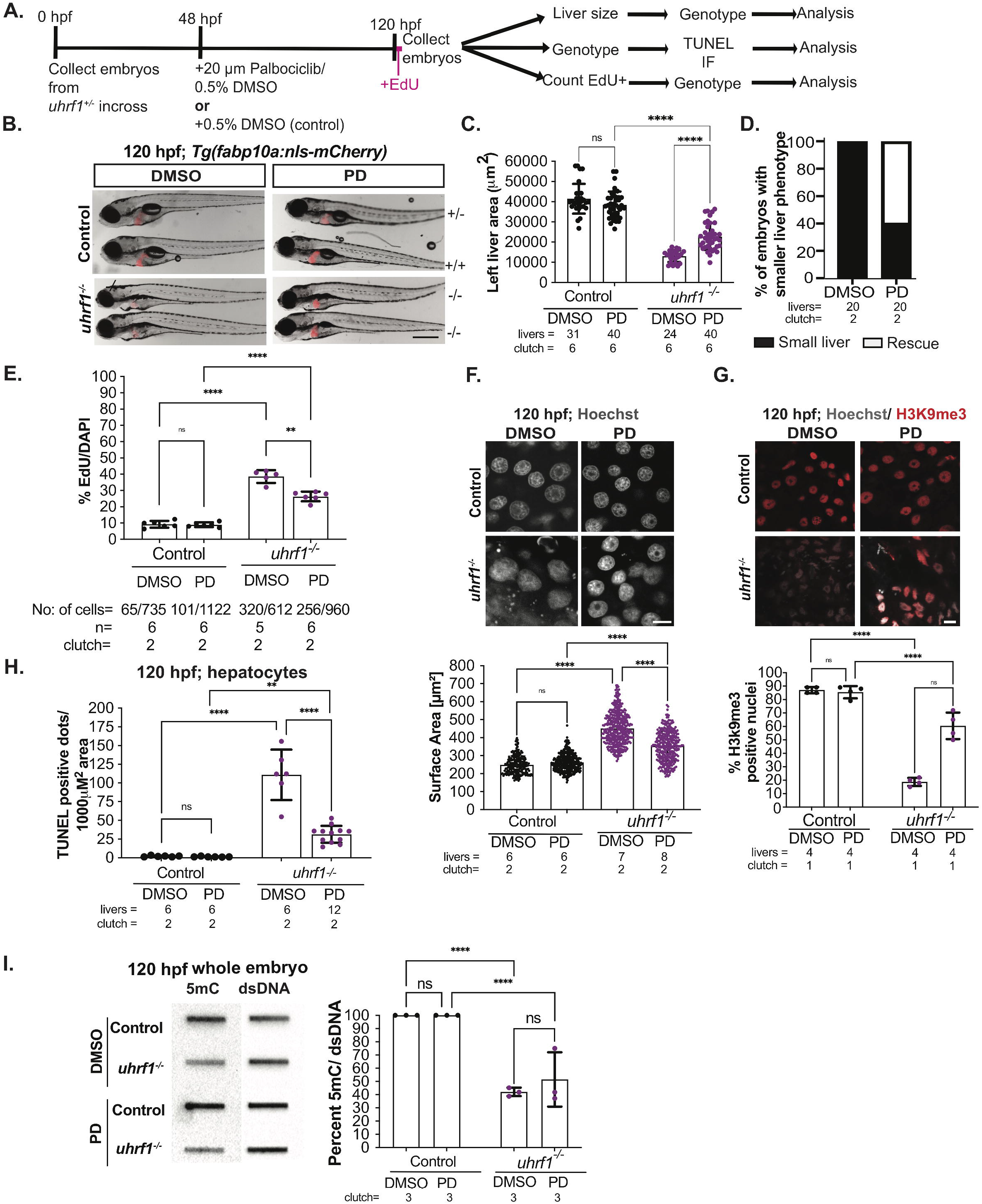
Cdk4/6 inhibition by Palbociclib (PD) rescues the *uhrf1* mutant liver phenotype. (A) Treatment scheme for inhibitor Palbociclib (PD). (B) Representative images of 120 hpf *uhrf1^-/-^* and control embryos after DMSO or PD treatment, liver indicated in red. (C) Quantification of the left liver lobe area of 120 hpf PD treated and untreated livers. (D) Liver phenotype rescue in 120 hpf *uhrf1*^-/-^ treated with PD. (E) quantification of 30 minute EdU incorporation at 120 hpf in DMSO and PD treated controls and *uhrf1*^-/-^. (F) Representative confocal image of 120 hpf Hoechst stained *uhrf1* control and *uhrf1*^-/-^ hepatocytes from DMSO or PD treatment and quantification of the nuclear surface area. (G) Representative immunofluorescent images of 120 hpf livers with heterochromatin marker H3K9me3 and quantification of H3K9me3 positive hepatocytes. (H) TUNEL assay was performed to detect cell death in 120 hpf in DMSO and PD treated controls and *uhrf1*^-/-^ and quantified. (I) Representative image of 5mC levels in *uhrf1*^-/-^ embryos and controls as measured by slot blot and quantification of the methylation levels from three clutches. Scale: 1000 µm in B, 50 µm in F,G, the number of samples and clutches indicated for each condition and represented as median with range. p-value *< 0.05, **< 0.005, *** < 0.0005 by unpaired Students t-test with adjustment for multiple comparisons.

At 120 hpf, *uhrf1* mutants are distinguished by smaller head, lower jaw deformity, microphthalmia, underdeveloped gut and small liver (Figure S2A; ^23, 25^). These phenotypes are fully penetrant at 120 hpf, so that on average, 25% of embryos resulting from an incross of *uhrf1^+/-^* adults display all of these phenotypes (Figure S2B), but only 7% of embryos produced from an incross of *uhrf1^+/-^* adults treated with 20 µM Palbociclib have all mutant phenotypes (Figure 3B). In treated clutches, 18% of embryos showed a partial mutant phenotype, with restoration of the lower jaw defect, increase in liver size, or both (Figure S2B-C). When these embryos were genotyped, we uncovered the expected genotype-phenotype relationship in 100% of embryos in DMSO treated clutches, so that all embryos displaying the mutant phenotypes were *uhrf1^-/-^* and all embryos that lacked these phenotypes were either *uhrf1^+/+^*or ^+/-^. However, in Palbociclib treated clutches, embryos with a partial phenotype and even a few scored as normal were *uhrf1^-/-^* (Figure S2C-D).

Palbociclib had no significant effect on liver size in WT embryos, but significantly increased the size of the liver (Figure 3B-D) and eye (Figure S2E) in *uhrf1* mutants. The liver size of nearly all *uhrf1* mutants treated with Palbociclib exceeded the average liver size in DMSO treated mutants, and 20% of treated mutants had a liver size that was within the range of WT embryos (Figure 3C). Strikingly, Palbociclib also increased survival of *uhrf1* mutants at 9 dpf (Figure S2F).

To differentiate aspects of the *uhrf1* mutant phenotype that were a direct consequence of Uhrf1 loss compared to those that were mediated by Cdk4/6 activation, we assessed the effects of Palbociclib on the cellular aspects of hepatic phenotypes: EdU incorporation, nuclear size and shape and cell death. These parameters were unchanged by Palbociclib treatment of control embryos, but were significantly rescued in *uhrf1* mutants. The number of hepatocytes with EdU incorporation at 120 hpf was reduced from 40% to 28% in untreated compared to Palbociclib treated *uhrf1* mutants (Figure 3E). This was accompanied by a dramatic change in the nuclear morphology: DMSO and Palbociclib treated control hepatocyte nuclei displayed clear nuclear rims and distinct heterochromatin foci, with prominent H3K9me3 staining, whereas *uhrf1* mutant nuclei were large and oblong, with significant nuclear debris (Figures 3F). The nuclear size, shape, DNA patterning and H3K9me3 staining in *uhrf1* mutant hepatocytes were nearly completely restored to the WT phenotype by Palbociclib treatment (Figure 3F-G). Finally, the amount of cell death in *uhrf1* mutant livers was significantly reduced by Palbociclib (Figure 3H). Therefore, Cdk4/6 activity is required for both the developmental and cellular phenotypes caused by Uhrf1 loss.

Since Uhrf1 is a required component of the maintenance methylation machinery ^8, 9^, DNA hypomethylation is a direct consequence of Uhrf1 loss. Assessment of bulk DNA methylation levels confirmed this, as both DMSO and Palbociclib treated *uhrf1* mutants had significant DNA hypomethylation (Figure 3I). Since Palbociclib treatment rescued most of the developmental and cellular defects in *uhrf1* mutants despite persistent DNA hypomethylation, we conclude that the developmental and cellular phenotypes caused by Uhrf1 loss is due to persistent activation of Cdk4/6 and is not a direct consequence of DNA hypomethylation.

### Cdk4/6 inhibition blocks cell cycle and Tnfa target genes activation in *uhrf1* mutant livers but does not suppress TE activation or the interferon response

We hypothesized that the cellular and developmental phenotypic rescue of *uhrf1* mutants by Palbociclib was attributed to downregulation of the G1/S transition and S-phase promoting genes. In contrast, we expected that activation of transposable elements (TEs) and the ensuing interferon response ^42, 45^ would not be affected by Palbociclib, since these are directly mediated by DNA methylation loss in *uhrf1* mutants.

We assessed this using bulk RNAseq analysis of the livers of 120 hpf *uhrf1* mutants and their phenotypically normal siblings (control) treated with DMSO or Palbociclib. Principal component analysis showed that controls cluster together regardless of treatment, while mutants treated with Palbociclib were closer to control samples (Figure 4A). Palbociclib treatment of control larvae resulted in modest downregulation of genes that play roles in hepatocyte function and metabolism, but had no effect on genes that regulate the cell cycle (Figure S3A-B; Table S4, S5). In contrast, Palbociclib dramatically changed the hepatic transcriptome in *uhrf1* mutants, reducing the number of DEGs from 2,272 in DMSO treated mutants to 650 in the liver of *uhrf1* mutants treated with Palbociclib, 560 DEGs common to both (Figure 4B, Table S6). While the common DEGs showed a very high correlation in their expression (r=0.974), the genes that were unique to each dataset genes were less correlated (r= 0.666; Figure S3C). In contrast, the upregulation of transposable elements (TEs) in *uhrf1* mutant livers was not suppressed by Palbociclib (Figure S3D).

**Figure 4.**
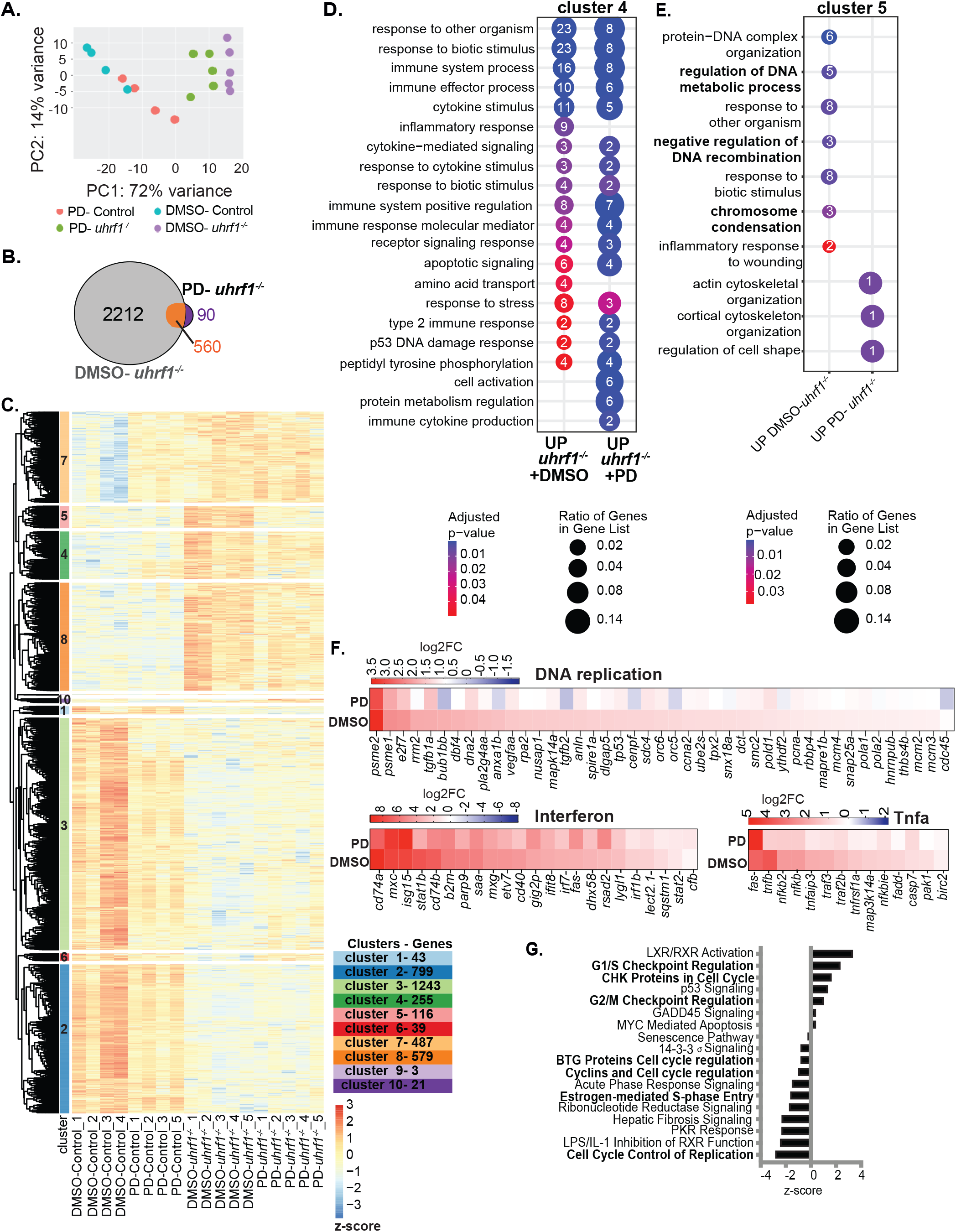
Cdk4/6 inhibition by Palbociclib (PD) rescues abnormal cell cycle gene expression in *uhrf1* mutants. (A) PCA analysis of RNA Seq from 120 hpf Palbociclib (PD) treated and DMSO treated *uhrf1*^-/-^ and control livers. (B) Venn diagram of differential expressed genes (DEGs) (cutoff *p_adj_* < 0.05) between *uhrf1*^-/-^ and their control siblings treated with DMSO and PD. (C) Unsupervised clustering heatmap of the expression profile of 120 hpf DMSO and PD treated *uhrf1*^-/-^ liver RNA-Seq with z-scores based on rows. Rows are divided into 10 clusters calculated on the hierarchical clustering of dendrogram (Euclidean distance). (D) REVIGO gene ontology of cluster 4 and (E) and cluster 5. (F) Key cell cycle genes, immune and *tnfa* response gene expression between PD and DMSO treated *uhrf1*^-/-^ plotted as a heatmap with log2fold change. (G) IPA of the cell cycle genes differentially expressed between 120 hpf *uhrf1*^-/-^ PD and DMSO livers, cell cycle checkpoint regulation pathways indicated in bold. p-value *< 0.05, **< 0.005, *** < 0.0005 by unpaired Students t-test with adjustment for multiple comparisons and represented as median with range.

We predicted that Cdk4/6 inhibition in *uhrf1* mutants would reduce the induction of E2F targets, but would have no effect on the interferon response as this is caused by TE activation. Unsupervised clustering of the unified set of DEGs from both DMSO and Palbociclib treated mutants showed that most of the upregulated genes in DMSO treated *uhrf1* mutant livers were still upregulated, but to a lesser extent in Palbociclib treated mutants (Figure 4C, cluster 5), however, a subset of genes remain elevated in both samples (cluster 4 and 5) (Figure 4C, Table S7). GO analysis of these two clusters shows that DNA replication related genesets were predominant in cluster 5, and that immune pathways were predominant in cluster 4 (Figure 4D-E, Table S8). Specific analysis of key genes involved in the pre-replication phase of cell cycle, such as *ccna2*, the *orcs*, *mcms* and *cdc45* which is important for origin firing at the replication forks in association with *mcms* ^53, 54^, showed that they were highly induced in DMSO treated mutants, but expressed at significantly lower levels in the liver of Palbociclib treated *uhrf1* mutants (Figure 4F, Figure S3E and Table S9).

Interestingly, the immune genes which were all upregulated in *uhrf1* mutants showed a nuanced response to Palbociclib treatment: interferon response genes remained highly expressed while genes downstream of Tnfa were downregulated in the liver of *uhrf1* mutants exposed to Palbociclib (Figure 4F, Table S10). Since CDK6 positively regulates TNFa ^55^, we conclude that Tnfa induction in *uhrf1* mutants is attributed to Cdk4/6 activation, in contrast to interferon related genes which are due to TE activation.

We further investigated the effect of Palbociclib on the expression of cell cycle regulatory genes in *uhrf1* mutants using IPA. We first analyzed the genes activated in *uhrf1* mutants compared to controls treated with DMSO and found significant activation of genes regulating G1/S *ccnd1*, *ccne* and *cdk4/6,* which lead to upregulation of transcription factors *e2fs, hdac, ftdp1*, as well as activation of DNA damage sensors *atm, atr* and *tp53* (Figure S4A). qPCR analysis of several of these genes confirmed that Palbociclib downregulated their expression the liver of *uhrf1* mutants (Figure S4B). We next compared the gene expression patterns in *uhrf1* mutants treated with Palbociclib to mutants treated with DMSO, and identified 280 DEGs (Figure S4C; Table S11). IPA analysis of these DEGs showed that genes involved in cell cycle checkpoints were upregulated and genes involved in cell cycle control and S-phase were downregulated in *uhrf1* mutant livers treated with Palbociclib (Figure 4G) but that Palbociclib had no effect on *atm/atr* and *tp53* induction in *uhrf1* mutants (Figure 4G, Figure S4D, Table S12), suggesting that DNA damage persists even when DNA replication is suppressed.

Together, these data indicate that Cdk4/6 activation does not affect DNA methylation, TE induction, interferon response and DNA damage caused by Uhrf1 loss, but is required for the downregulation of G1/S checkpoints, E2F targets, DNA rereplication and cell death. Based on our previous finding that blocking Tnfa reduced cell death in *uhrf1* mutant livers but has no effect on the mutant phenotype or liver size at 120 hpf ^42^, combined with the current finding that Palbociclib reduced Tnfa activation, we deduce Palbociclib prevents cell death in *uhrf1* mutants by inactivation of Tnfa.

### Cdk4/6 is activated by *dnmt1* mutation

To understand if the cellular phenotypes that cause hepatic outgrowth failure in *uhrf1* mutants are attributed to DNA hypomethylation, we assessed larvae with a loss of function mutation in *dnmt1* ^24^. These have a strikingly similar phenotype to *uhrf1* mutants, including DNA hypomethylation, large and disorganized hepatocyte nuclei and a small liver ^25, 26, 41, 42, 56^. We asked whether the cell cycle related geneset that we found to be differentially expressed in *uhrf1* mutant livers (the unified set of all cell cycle genes in Figure 2B) were also differentially expressed in bulk RNAseq data from the liver of 120 hpf *dnmt1* mutants ^42^. Similar to *uhrf1* mutants, many of the genes required for DNA replication (*pcna, orcs, mcms*, *cdc45* and *pold2*) were upregulated in *dnmt1* mutant livers (Figure 5A). Surprisingly, however there were a lower number of hepatocytes that incorporate EdU in *dnmt1* mutants at both 96 hpf and 120 hpf (Figure 5B). Dnmt1 loss prevents DNA replication, consistent with a requirement for Dnmt1 at the replication fork ^11, 12^. Previous analysis showed that *uhrf1* and *dnmt1* mutant livers have a similar pattern whereby genes required for DNA replication were upregulated ^25, 42^. IPA provided a more detailed analysis, while the Atm and Tp53 mediated DNA damage responses were upregulated in both samples, some genes that regulate S-phase (*chek1, chek2, tlk1/2, cdc25),* M phase progression (*cdk1, cyclin B*) and Atr mediated DNA damage response signaling were upregulated only in *uhrf1* mutant livers (Figure S5). Thus, while both mutants have a global upregulation of cell cycle promoting genes and the Atm mediated response to double stranded breaks, *uhrf1* loss specifically causes the response to single stranded breaks.

**Figure 5.**
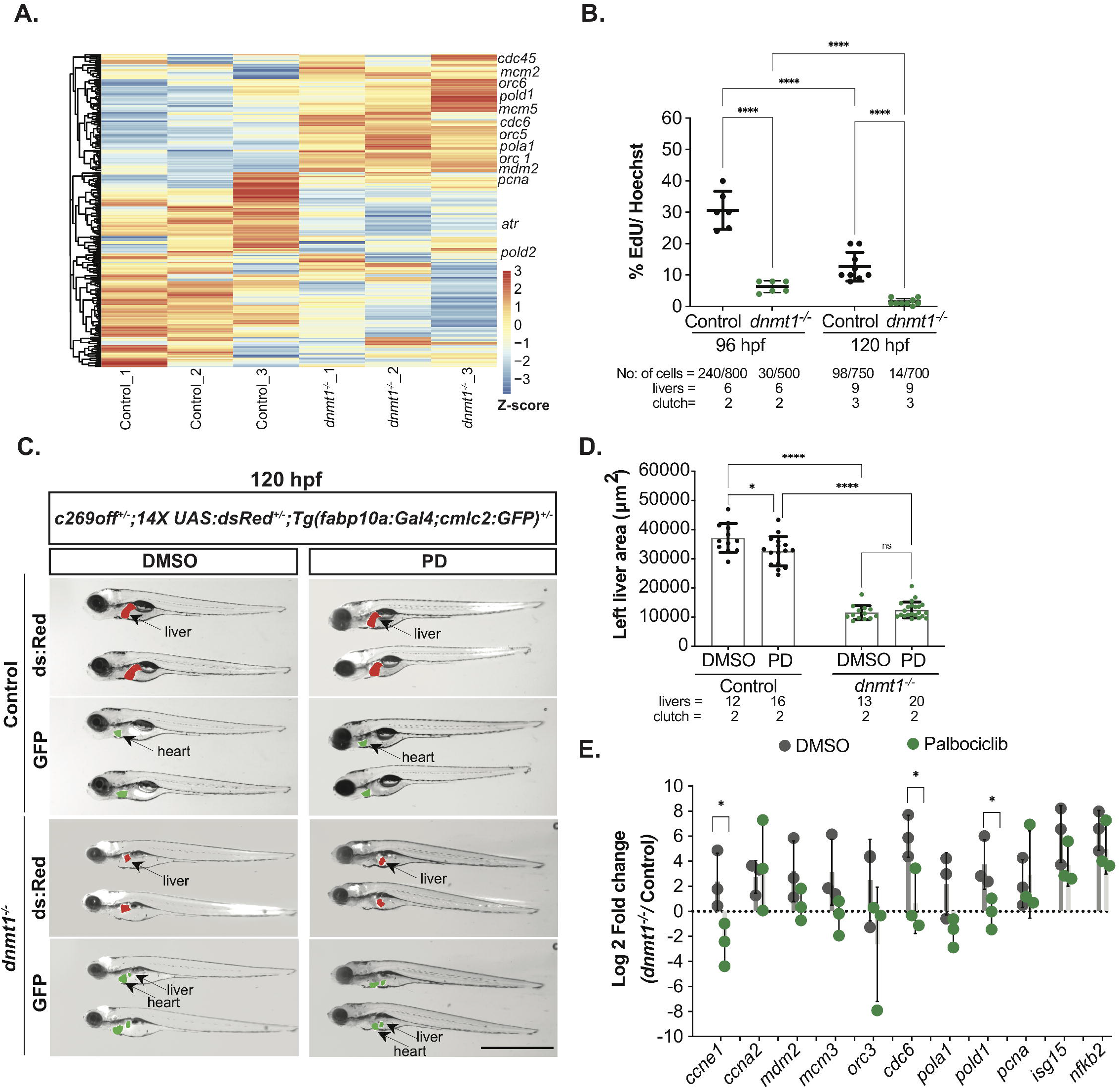
Loss of *uhrf1* leads to Cdk4/6 mediated Dnmt1 protein stabilization and smaller liver size. (A) Unsupervised clustering heatmap of all the cell cycle genes of 120 hpf *dnmt1^-/-^* liver RNA-Seq with z-scores based on rows, important S-phase genes are indicated. (B) EdU quantification (pulse 30 minute) of 96 hpf and 120 hpf control and *dnmt1^-/-^* hepatocytes. (C) Representative images of 120 hpf *dnmt1^-/-^* and controls with *c269off+/-; 14X UAS:dsRed+/-; Tg(fabp10a:Gal4;cmlc2:GFP)+/-* treated with DMSO or Palbociclib (PD), showing the liver (dsRed), the heart (GFP) as a marker of transgenesis and in green, the liver with hypomethylation in mutants only. (D) Quantification of the left liver lobe area of *dnmt1^-/-^* and controls, treated with PD or DMSO. Scale: 50 µm, the number of samples and clutches indicated for each condition. (E) QPCR of key cell cycle genes and immune genes in *dnmt1^-/-^* livers. Scale: 1000 µm in C, the number of samples and clutches indicated for each condition. p-value *< 0.05, **< 0.005, *** < 0.0005 by unpaired Students t-test with adjustment for multiple comparisons and represented as median with range.

If Cdk4/6 is upstream of cell cycle activation in *uhrf1* and *dnmt1* mutants, treatment of *dnmt1* mutants with Palbociclib should downregulate these genes. Palbociclib had no significant effect on the developmental phenotype or liver size in *dnmt1* mutants (Figure 5C-D). While there was variability in the induction of cell cycle genes in *dnmt1* mutant livers, treatment with Palbociclib decreased the expression of assessed cell cycle genes, with *ccne1, cdc6* and *pold1* reaching significance. As in *uhrf1* mutants, Palbociclib had no effect on interferon response gene expression in *dnmt1* mutant livers (Figure 5E). Thus, while *dnmt1* and *uhrf1* loss activate Cdk4/6 and E2F target genes, derepress TEs and induce an interferon response, these responses are independent. Induction of cell cycle genes requires Cdk4/6 signaling while TE activation and the interferon response do not. Moreover, the finding that *dnmt1* mutants are not rescued by Palbociclib reflects the fact that Dnmt1 deficient cells cannot undergo DNA replication, likely due to the requirement of Dnmt1 at the replication fork ^12, 57, 58^.

### DNA replication in *uhrf1* mutant hepatocytes requires *dnmt1*

We examined the relationship between Uhrf1 deficiency, DNA replication and Dnmt1. Western blotting of whole embryos from control and *uhrf1* mutants ^59^ showed a significant increase in Dnmt1 levels starting at 80 hpf in mutants (Figure 6A), suggesting that Dnmt1 upregulation is an early molecular consequence of *uhrf1* mutation. Cdk4/6 activity has been shown to stabilize Dnmt1 ^60^ and thus one mechanism by which Palbociclib could reduce DNA rereplication in *uhrf1* mutants is by reducing Dnmt1 levels. To test this, we assessed Dnmt1 protein levels in 120 hpf hepatocytes using immunofluorescence. Dnmt1 was not detected in controls, but there was strong expression in 70% of *uhrf1* mutant hepatocytes, and this was depleted in *uhrf1* mutants treated with Palbociclib treatment (Figure 6B). Therefore, Dnmt1 protein stabilization in *uhrf1* deficient hepatocytes requires Cdk4/6. This could either be a direct effect of Cdk4/6 on Dnmt1 stability or could be indirect via induction of dnmt1 transcriptional activation by E2F.

**Figure 6.**
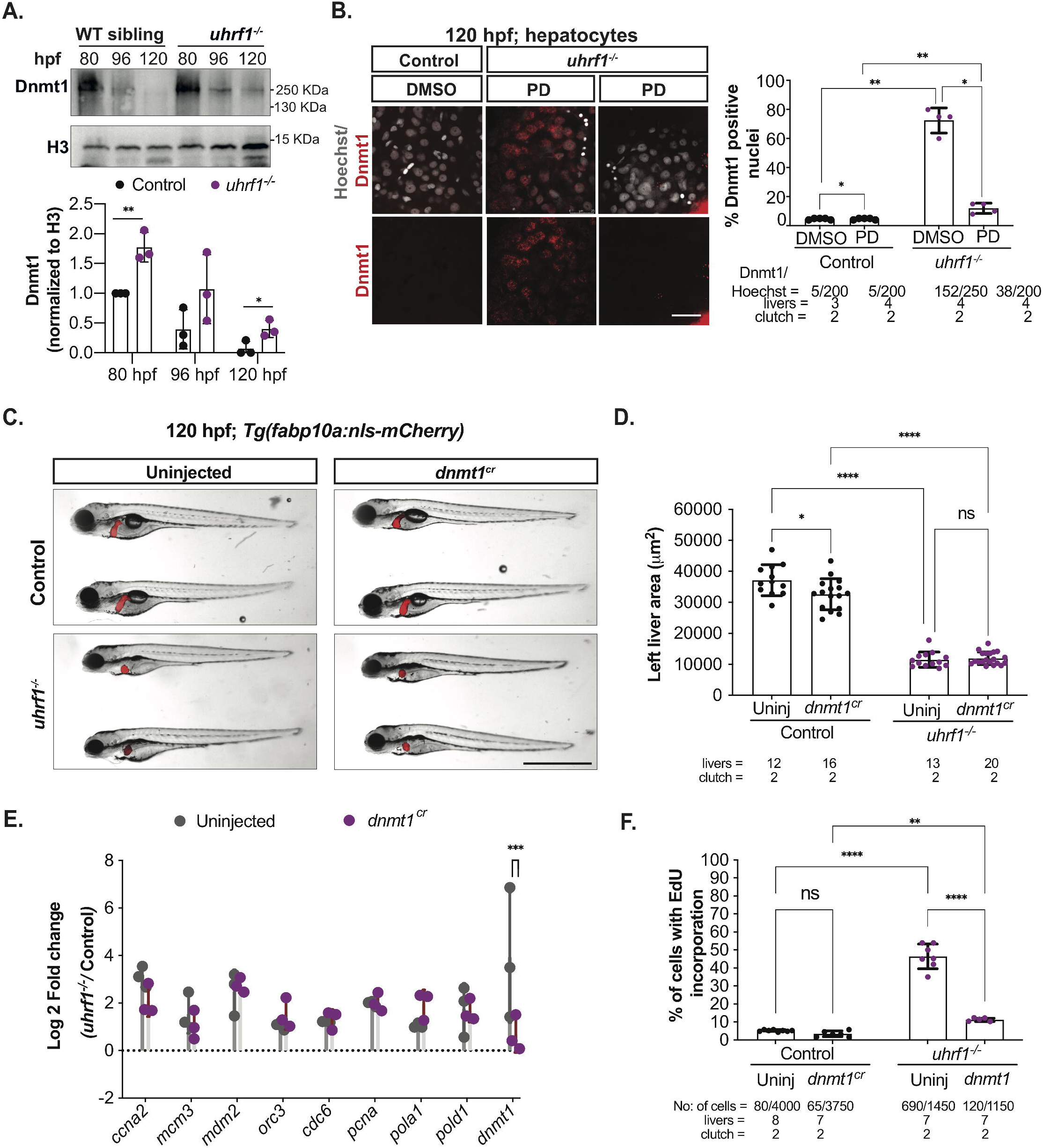
*dnmt1* mutation rescues DNA replication defect in *uhrf1* mutant hepatocytes. (A) Western blot of 80, 96 and 120 hpf *uhrf1^-/-^* and control siblings, levels of Dnmt1 are normalized on level of H3 for each sample and plotted. (B) Representative immunofluorescent images with Dnmt1 staining of 120 hpf control and *uhrf1^-/-^* livers treated with DMSO or PD and Dnmt1 positive cell quantification. (C) Representative images of 120 hpf *uhrf1^-/-^* and control embryos uninjected or injected with *dnmt1^cr^.* (D) Quantification of the left liver lobe area of 120 hpf *uhrf1^-/-^* and controls uninjected or injected with *dnmt1^cr^*. (E) QPCR of key cell cycle genes in 120 hpf livers of *uhrf1^-/-^* and controls uninjected or injected with *dnmt1^cr^*. (F) Representative images of 30 minute EdU incorporation at 120 hpf in uninjected and *dnmt1^c^*^r^ injected controls and *uhrf1^-/-^* livers. Scale: 50 µm in B, 1000 µm in C, the number of samples and clutches indicated for each condition p-value *< 0.05, **< 0.005, *** < 0.0005 by unpaired Students t-test with adjustment for multiple comparisons and represented as median with range.

To determine if Dnmt1 was required for the EdU incorporation phenotype in *uhrf1* mutant hepatocytes, we used CRISPR-Cas9 to target *dnmt1*. Efficacy was verified using an *in vivo* assay for DNA methylation in which the UAS:GFP cassette is silenced by DNA methylation, and GFP expression occurs in response to Gal4 when methylation is lost on the UAS promoter (*Tg(c269off^+/-^; 14X UAS:dsRed)* ^42, 61^. All *Tg(c269off^+/-^; 14X UAS:dsRed^+/-^);dnmt1^cr^* larvae had GFP expression in the head at 120 hpf (Figure S6A), but had a mild phenotype compared to *dnmt1* mutants (Figure 5C). Crispr deletion of *dnmt1* did not change liver size of *uhrf1* mutants (Figure 6C-D) or the volume or shape of the *uhrf1* mutant hepatocyte nuclei (Figure S6B-D). *dnmt1* deletion did not rescue aberrant expression of cell cycle genes in *uhrf1* mutant livers (Figure 6E) but did significantly reduce EdU incorporation in hepatocytes compared to uninjected *uhrf1^-/-^* (Figure 6F). We conclude that Dnmt1 is required for DNA replication in *uhrf1* mutant hepatocytes.

### Atr is required for DNA replication and hepatic outgrowth failure in *uhrf1* **mutants**

ATR responds to DNA replication stress by stabilizing replication forks, delaying or blocking the progress of the cell cycle, promoting DNA repair ^62, 63^. In response to replication stress, ATR activates dormant origins while repressing firing within those that are not yet activated ^64^. We found Atr to be specifically activated in *uhrf1* mutants, but not *dnmt1* mutants, suggesting that the DNA replication phenotype in *uhrf1* mutants, which is absent from *dnmt1* mutants, causes replication stress mediated Atr activation. To test this, we established a treatment protocol using the highly selective and potent ATR inhibitor VE-821 ^65^ during zebrafish development. While treatment from 48-120 hpf with 10 µm VE-821 had no effect control larvae and had minimal effects on the overall *uhrf1* mutant phenotype, it significantly increased liver size (Figure 7A-B). Moreover, EdU incorporation in hepatocytes was reduced from 36% to 24% in untreated and VE-821 treated *uhrf1* mutants, respectively (Figure 7C-D). This suggests that DNA replication in uhrf1 mutants is dependent on Atr.

**Figure 7.**
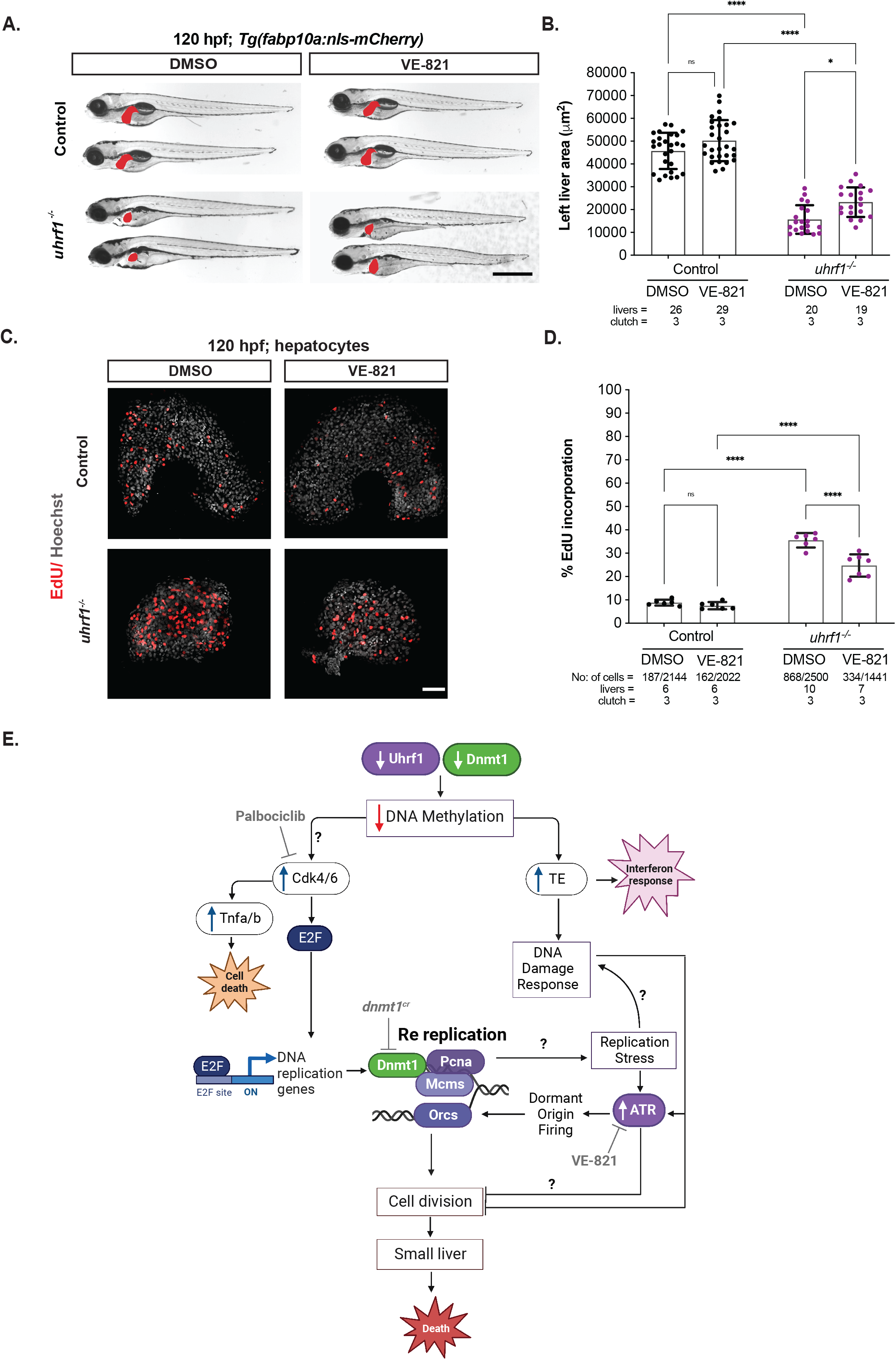
Atr inhibition rescues liver size and DNA rereplication in *uhrf1* mutant hepatocytes. (A) Representative images of 120 hpf *uhrf1^-/-^* and control larvae after DMSO or VE-821 treatment, liver indicated in red. (B) Quantification of the left liver lobe area of 120 hpf VE-821 treated and untreated embryos from *uhrf1^-/-^* and control larvae. (C) Representative Z-stack projection of 120 hpf EdU and Hoechst stained control and *uhrf1*^-/-^ livers from DMSO or VE-821 treated larvae. (D) Quantification of 30 minute EdU incorporation at 120 hpf in DMSO and VE-821 treated controls and *uhrf1*^-/-^ livers. Scale: 50 µm in A, 50 µm in C, the number of samples and clutches indicated for each condition p-value *< 0.05, **< 0.005, *** < 0.0005 by unpaired Students t-test with adjustment for multiple comparisons. Data represented as median with range. (E) Model of the relationship between *uhrf1* mutation, DNA methylation, cell cycle regulation, DNA replication and small liver. Uhrf1 and Dnmt1 loss cause DNA hypomethylation, upregulation of TEs and inflammatory response. Cdk4/6 activation upon DNA hypomethylation causes Tnfa/b activation, leading to cell death in *uhrf1* deficient livers. Cdk4/6 dependent activation of DNA replication genes leads to DNA rereplication, replication stress and Atr activation. Both TE activation and replication stress trigger DNA damage response to inhibit cell division, restricting organ outgrowth and causing lethality in *uhrf1* mutants.

## Discussion

We report that Cdk4/6 activation is the mechanism accounting for the unusual cell cycle defect caused by *uhrf1* mutation in zebrafish and present a model that illustrates the relationship between DNA methylation, cell cycle regulation, DNA replication and development (Figure 7E). The striking rescue of the cellular and developmental defects in Uhrf1 deficient zebrafish embryos by Cdk4/6 inhibition suggests that S-phase entry causes all the downstream phenotypes. The data presented here, combined with our finding that reducing cell death does not rescue liver size ^42^, indicates that the cell cycle defect is the main mechanism of hepatic outgrowth failure. Surprisingly, the cellular and developmental phenotypes resolve, despite persistent DNA hypomethylation, TE activation and an interferon response. This finding differentiates between the direct targets of Uhrf1 – i.e. DNA hypomethylation – and those that are secondary consequences – i.e. DNA rereplication. These data also show that loss of DNA methylation and TE activation *per se* do not cause the developmental defects that occur in *uhrf1* mutants. Rather it is the downstream consequences of DNA hypomethylation which are responsible.

The finding of similar Cdk4/6 dependent gene expression in *dnmt1* and *uhrf1* mutants suggests that DNA hypomethylation is the mechanism for Cdk/4/6 activation, although the mechanism of Cdk4/6 activation is not clear. We previously published that *uhrf1* mutant hepatocytes have both an abnormal DNA replication pattern and widespread cell death ^23, 25^. Importantly, the role of Uhrf1 in regulating the hepatic cell cycle is conserved in the developing mouse liver. We demonstrate that both *uhrf1* and *dnmt1* mutants activate of cell cycle genes in a Cdk4/6 dependent manner, but while *uhrf1* mutant hepatocytes undergo DNA rereplication, *dnmt1* mutant hepatocytes do not. Since DNA replication in *uhrf1* mutant hepatocytes was rescued by *dnmt1* depletion, we conclude that either the *uhrf1* mutant phenotype could be partly attributed to elevated Dnmt1 levels or, more likely, it is due to the requirement for Dnmt1 for a functional replication fork ^57, 58^. Hence, Dnmt1 is both upstream and downstream of Cdk4/6 in this model. The finding that Atr activation is required for DNA replication and small liver size in *uhrf1* mutant hepatocytes suggests Atr signaling activates DNA replication and prevents cell cycle progression. Taken together, these results provide mechanistic insight into cell cycle regulation during development and point to a functional relationship between epigenetic regulation and cell cycle control.

Uhrf1 loss has been implicated in the mis-regulation of both G1 and G2/M phases of the cell cycle ^18, 28, 66^ and here we provide evidence for a functional role in regulating DNA replication. There are several possible mechanisms underlying this phenotype. The first, discussed above, is that DNA replication stress activates Atr to prevent cell cycle progression while promoting dormant origin firing (Figure 7E). During normal DNA replication, fork speed has to be moderated for the fidelity of normal cell cycle, with an increased fork speed triggering replication stress and DNA damage response ^67, 68^. However during DNA rereplication, DNA synthesis was reported to progress slowly, and repeatedly utilize the same main origins ^69^. In contrast, unscheduled firing of dormant origins can lead to increased and uncoordinated DNA synthesis ^3, 70^. While the experiments required to assess fork speed is challenging in whole embryos, the Atr inhibition data in *uhrf1* mutants supports the hypothesis that the DNA rereplication in *uhrf1* deficient hepatocytes is caused by Atr dependent firing of multiple dormant origins, not a repeated firing of the same origins. The question of why origins of replication can be fired at all if there is a checkpoint activated by ATR can be answered by one study where ATR/ Chk1 pathway activation in response to replication stress caused dormant origin firing within active replication clusters ^71–73^. Hence, in *uhrf1* mutants, Atr activation could be activated by two mechanisms: either directly by Cdk4/6 and/or as a downstream effect of replicative stress brought on by Cdk4/6 activation. Further investigation between the CDK4/6 pathway and the players in the ATR pathway will provide better insight linking cell cycle checkpoint, replication stress and dormant origin firing leading to DNA re replication in *uhrf1* deficient hepatocytes.

Another possible mechanism of the DNA replication defect in *uhrf1* mutants could involve a defect in regulating which origins fire due to a failure to degrade replication licensing factor Cdt1, which is required to prevent reuse of replication origins. Since the levels of Cdt1 protein are regulated by the Cul4–Ddb1–Cdt2 E3 ubiquitin ligase complex ^74^, it is feasible that Cdt1 stabilization due to nuclear Cyclin D1/CDK4, as shown in other models ^75^, could result in repeated origin use. Further, Cyclin D1/CDK4 during S-phase specifically represses transcription of *CUL4,* resulting in CDT1 stabilization ^76^. Our finding that *uhrf1* loss induced overexpression of *ccnd1* and *cdk4* and downregulation of *cul4* suggests that Ctd1 levels may persist and promote DNA rereplication ^77^. Further biochemical analysis is required to evaluate this mechanism.

The most straightforward conclusion to explain how Palbociclib prevents DNA replication in *uhrf1* mutants is through reinstatement of Rb mediated suppression of E2f. Another possibility is that Cdk4/6 inhibition depletes Dnmt1, which we have shown is required for DNA re replication. Uhrf1 targets Dnmt1 for degradation ^78–82^ and CDK4 phosphorylates DNMT1, preventing its degradation and therefore Palbociclib treatment reduces DNMT1 protein levels ^83^. Our data suggest that in *uhrf1* mutant embryos, Dnmt1 accumulation is attributed to increased Cdk4/6 activity, as inhibition of Cdk4/6 caused significant reduction in Dnmt1 levels. However, it is also possible that this effect was mediated through decreased E2f mediated *dnmt1* transcription. We conclude that Dnmt1 levels are permissive to the unscheduled replication phenotype in *uhrf1* mutant hepatocytes.

We were intrigued by the finding that Palbociclib blocked cell death in *uhrf1* mutant hepatocytes. In other studies, Palbociclib has been shown to activate NF-κB and TNF-induced gene expression in mammalian cells ^84^. Given our previous finding that Tnfa depletion in *uhrf1* mutants prevented hepatocyte cell death but did not rescue liver size^42^, we attribute the rescue of cell death in Palbociclib treated *uhrf1* mutants to the downregulation of Tnfa signaling, and conclude that cell death is not caused by cell cycle defects. These data also show that the small liver phenotype in this context is not caused by cell death, but rather is due to failure of hepatocyte proliferation.

Our data point to DNA hypomethylation as an upstream signal that leads to Cdk4/6 activation in hepatocytes. How this occurs is not known, but other studies suggest that the p16^INK4a^ inhibitor of CDK4/6 could be suppressed through DNA methylation ^31, 85^, and inhibiting p16^INK4a^ leads to increased cell cycle progression and increased DNMT1 expression ^31^. While it would be surprising if *uhrf1* and *dnmt1* mutation caused hypermethylation of the p16^INK4a^ regulatory sequences or any other Cdk4/6 inhibitor, it is possible that loss of methylation causes redistribution of another repressive epigenetic marks to the p16^INK4a^ regulatory regions, as we described in other contexts ^44^. Alternatively, loss of DNA methylation could derepress Cyclin D, leading to perpetual Cdk4/6 activation.

In summary, this work provides the first insight into two distinct functions of Uhrf1 during liver development. Here we show that Uhrf1 loss induces Cdk4/6, E2f and all the genes that are required for S-phase. This is paradoxical, as UHRF1 is overexpressed in nearly all types of solid tumors ^86^, including HCC ^78, 87^, Palbociclib effectively suppressed cell proliferation in tumor growth in preclinical models of HCC by promoting a reversible cell cycle arrest ^52^. Therefore, it is possible that in HCC, UHRF1 overexpression can act as a dominant negative to activate CDK4/6 and promote cell proliferation, and future studies may investigate whether Palbociclib functions as a therapeutic by counteracting the oncogenic effects of UHRF1.

## Materials and Methods

### Zebrafish Husbandry and Genotyping

Adult zebrafish were maintained in a circulating aquaculture system on a 14:10 h light: dark cycle at 28 °C. The *uhrf1*^hi2^^72^ allele ^88^ was maintained by genotyping adults to identify heterozygous carriers as described ^88^. Homozygous mutant embryos were generated by crossing heterozygous adults and were identified based on distinctive phenotypes as described ^23, 25, 26^ or by genotyping individual embryos as described ^23^. *uhrf1^hi272+/−^*adults were genotyped by PCR as described ^23^. *Tg(fabp10a: nls-mcherry)* in *uhrf1^hi272+/−^* and *Tg(fabp10a: CAAX-EGFP)* in *uhrf1^hi272+/−^* background was used for all experiments for ease of liver detection ^78^. Homozygous *uhrf1^hi2^*^72^*^-/−^*mutant larvae are hereafter called *uhrf1^−/−^*. Homozygous *dnmt1^s9^*^04^*^-/-^* mutants (*dnmt1^-/-^)* were generated by crossing *dnmt1^s904+/−^* adults in *dnmt1^s904+/−^; Tg(c269*^o*ff*^*; 10XUAS:dsRed); (fabp10a: Gal4; cmlc2: GFP)* background to monitor for DNA methylation in the liver as described ^42^. All protocols were approved by the NYU Abu Dhabi Institutional Animal Care and Use Committee (IACUC).

### Mice Husbandry and Usage

Mice maintenance and experiments were approved by the New York University Abu Dhabi Animal Care and Use Committee. Temperature, humidity, and light:dark cycles were controlled and mice were fed food and water ad libitum. Mice with hepatocyte-specific deletion of the Uhrf1 gene (*Uhrf1^fl/fl^;Alb-Cre*; referred to as *Uhrf1^HepKO^* mice) were generated by crossing mice homozygous for floxed *Uhrf1* (*Uhrf1^fl/fl^*) with mice expressing the Cre recombinase under the liver-specific albumin promoter back-crossed onto the *Uhrf1^fl/fl^* background (*Alb-cre^Tg/+^; Uhrf1^fl/+^*) described previously ^44^. P4 mouse embryos were sacrificed and their livers were collected, flash frozen in liquid nitrogen, and stored at –80 °C for subsequent analysis.

### FACS Analysis of the Cell Cycle

For FACS, 20 *uhrf1^-/-^; Tg(fabp10a: nls-mcherry)* and 20 *Tg(fabp10a: nls-mcherry)* sibling larvae were collected and used for single cells suspension. Single cell suspension was prepared by incubating larvae at 37 °C for 20 minutes in Trypsin-EDTA 0.05% (Thermo Fisher). After 20 minutes 10% FBS was added to stop the activity of the trypsin and cells were centrifuged at 2000 g for 7 minutes at 4 °C followed by 2 washes in PBS. Cells were resuspended in PBS and fixed with ice cold ethanol overnight. Cells were staining in PBS containing 10 μg/ml Hoechst (Thermo Fisher) for 10 minutes in dark and washed twice with PBS. Cells were finally resuspended in PBS containing 1% BSA (Sigma Aldrich) and 10 μg/ml RNaseA (Invitrogen) and acquired at FACS Aria II (BD Bioscience). Hepatocytes were selected by gating for mCherry^+^ population, and Hoechst levels recorded for analysis. Analysis was performed with FlowJo 10 and plotted in GraphPad Prism 9.

### CRISPR/Cas9 Generation and T7 Endonuclease Assay

4 sgRNAs for *dnmt1 was* designed by using ChopChop (https://chopchop.cbu.uib.no/). sgRNA for *slc45a2* (gene involved in pigmentation) were previously designed and validated ^40, 89^. Genotyping primers were designed by Primer3 (https://bioinfo.ut.ee/primer3-0.4.0/) and validated in USCS Genome Browser (https://genome.ucsc.edu/cgi-bin/hgPcr). sgRNAs were produced by sgRNA IVT kit (Takara Bio) by following the manufacturer’s instructions and RNA was isolated by Trizol (Invitrogen). sgRNAs were quantified by Qubit RNA BR kit and diluted at 50 ng/μl and stored as single use aliquots. The efficiency of the sgRNAs were assessed by injecting all 4 *dnmt1* sgRNAs into WT embryos with equal volume of previously diluted nls-Cas9 protein (IDT; 0.5 μl of nls-Cas9 added with 9.5 μl of 20 mM HEPES; 150 mM KCI, pH 7.5) and sgRNAs, incubated at 37 ^◦^C for 5 minutes and then 1 nl was injected in 1–2 cells stage embryos. At 24–72 hpf, 12–16 sgRNA injected embryos were individually collected and genomic DNA was extracted by heat shock denaturation in 50 mM NaOH (95 ^◦^C for 20 minutes). For each embryo, PCR was performed on genomic DNA by using Q5 High-Fidelity Taq Polymerase (New England Biolabs) followed by T7 endonuclease I assay (New England Biolabs) to detect indel mutations. For T7 endonuclease I assay, 10 μl of PCR product was incubated with 0.5 μl of T7e1 enzyme (New England Biolabs) for 30 minutes at 37^◦^C. Digested and undigested fragments were run in parallel in 2% agarose gel to assess the presence of indels. Efficiency was calculated as the number of embryos that show a positive result based on T7e1 assay divided by the total number of embryos assayed for the sgRNAs. Once established for efficiency, the pool of 4 *dnmt1* sgRNAs were injected into the 1 cell embryos generated by an incross of *uhrf1*^+/-^ adults as previously described. Not injected embryos or *slc45a2* crispants were used as control. The resulting F0 larvae were considered crispants. For each clutch, *uhrf1*^-/-^ mutants were divided from phenotypically WT siblings at 5 dpf based on morphological differences and T7e1 assay. sgRNA sequence information and genotyping primers are provided in Table S13.

### Inhibitor Treatment

The CDK4/6 inhibitor Palbociclib (PD-0332991, Sigma Aldrich) ^52^ was dissolved in DMSO (Sigma) to yield a 10 mM stock solution and stored at –80 °C. For the *in vivo experiments*, Palbociclib was dissolved in 0.5% DMSO embryo water at a concentration of 20 μM. 48 hpf dechorionated embryos in 6 well plates were treated either with Palbociclib in 0.5% DMSO or 0.5% DMSO alone and collected at 120 hpf for further analysis. The ATR inhibitor VE-821 (ab219506, Abcam) ^90^ was dissolved in DMSO (Sigma) to yield a 20 mM stock solution and stored at –80 °C. For the *in vivo experiments*, VE-821 was dissolved in embryo water at a concentration of 10 μM. 48 hpf dechorionated embryos were treated either with VE-821 or DMSO (1:2000 dilution) in embryo water and collected at 120 hpf for further analysis.

### Protein Lysates and Western Blotting

For western blot, whole larvae from *uhrf1^-/-^* and control siblings were collected at 80, 96 and 120 hpf in RIPA buffer (10mM Tris-HCl pH 8.0, 1mM EDTA, 0.5mM EGTA, 1% Triton X-100, 0.1% Sodium Deoxycholate, 0.1% SDS, 140mM NaCl supplemented with protease inhibitor cocktail (Machery-Nagel), sonicated using a hand sonicator (Hielscher Ultrasonics), and centrifuged at 14000 g for 15 minutes at 4°C to remove cell debris. Supernatant was collected and SDS-PAGE loading Leammli buffer (BioRad) was added and run on SDS-PAGE gel. Volume corresponding to 4-5 larvae was loaded on 10% acrylamide gels, transferred onto PVDF membranes, blocked with 5% w/v powdered skimmed milk (Sigma) in TBST (20 mM Tris–HCl, 150 mM NaCl, 0.1% v/v Tween 20, pH 8.0) for 1 hour at room temperature and incubated overnight in primary antibody (1:100 anti-Dnmt1 polyclonal, Santa Cruz, 1:1000 H3 polyclonal, Santa Cruz) diluted in blocking buffer at 4°C. The following day membranes were washed 3 times n TBTS for 5 minutes and incubated in secondary antibody (1:2500 anti-Rabbit HRP conjugated, Promega) diluted in blocking buffer for 1 hour at room temperature and washed 3 times with TBTS for 5 minutes at room temperature. Chemiluminescent signals using Pierce™ ECL Western Blotting Substrate (Thermo Fisher Scientific) or Clarity ECL substrate (BioRad) and imaged using Bio-Rad ChemiDoc. Immunoblot bands are quantified by densitometry using GelAnalyzer (http://www.gelanalyzer.com) and plotted using GraphPad Prism 9. 3 independent clutches were used for quantification. Levels of Dnmt1 are normalized on level of H3 for each sample and then for the level of Dnmt1 of 80 hpf wt looking larvae for that clutch.

### RNA and DNA Extraction

For each clutch, 10 to 20 livers were microdissected from *dnmt1^-/-^* and their controls or *uhrf1^-/-^* and their controls in Tg(fabp10a: CAAX-EGFP) background larvae to facilitate liver identification according to published protocols ^91^. RNA was extracted using Trizol (Invitrogen) following the manufacturer’s instructions with modifications. Briefly, during precipitation in isopropanol, 10 μg of Glycoblue (Thermo Fisher Scientific) was added, and precipitation performed overnight at −20 °C followed by 1 h centrifugation at 12,000× g at 4 °C. The resultant RNA pellet was resuspended in water and quantified by Qubit RNA High Sensitivity kit (Thermo Fisher). For genomic DNA extraction, each embryo or 10 livers were subjected to DNA extraction buffer (10 mM Tris-HCl pH9, 10 mM EDTA, 200 mM NaCl, 0.5% SDS, 200 μg/mL proteinase K) followed by DNA precipitation with isopropanol as previously described ^45^. DNA was resuspended in water and quantified by Qubit dsDNA High Sensitivity kit (Thermo Fisher).

### Slot Blot Analysis of 5-MeC

Slot blot was performed as previously described ^25^. Briefly, 2 ng of genomic DNA was denatured in 400 mM NaOH/10 mM EDTA and blotted onto nitrocellulose membrane in duplicate for dsDNA and 5-MeC using a slot blot apparatus. Membranes were incubated 1 h at 80 ^◦^C, blocked with 5% skim milk in TBST (37 mM NaCl, 20 mM Tris pH 7.5, 0.1% Tween 20), and incubated overnight at 4 ^◦^C in either anti-dsDNA (Abcam, 1:8000 in 2% BSA in TBST) or anti-5-methyl-cytosine (m5C– Aviva Biosystem clone 33D3, 1:2000 in 2% BSA in TBST). Membranes were washed in TBST and probed with anti-mouse HRP secondary antibody (Promega; 1:5000 in 2% BSA in TBST) for 1 h at room temperature followed by development in ECL (Thermo Fisher Scientific). ChemiDoc (BioRad) was used to detect and quantify the chemiluminescent signal. Gel Analyzer was used to measure the signals and ratio between 5-MeC and dsDNA was plotted for controls and mutants in each clutch using GraphPad Prism 9.

### cDNA Production and qPCR

RNA extracted from microdissected livers was retrotranscribed to cDNA using Qscript cDNA synthesis kit (Quanta Bio) following the manufacturer’s instructions. cDNA was diluted to 1 ng/μL, and 5 μL was used per reaction for qPCR using Maxima R SYBR green/ROX master mix (Thermo Fisher Scientific). *rplp0* was used to normalize expression levels by using the calculations for delta-Ct, and control siblings were used to calculate delta-delta-Ct (DDCt) as previously described ^92^. To determine changes in expression between control and experimental samples, the fold change was calculated and expressed as the log 2 value (L2FC) for display. All experiments were performed on samples from at least two independent clutches, with the number of biological replicates indicated for each experiment. Primer information is provided in Table S13.

### Left Lobe Liver Size Measurements

Liver size measurements were carried out based on the 2-dimensional area of the left liver lobe in 120 hpf larvae. Embryos from incross of *Tg(fabp10a: nls-mcherry)* in *uhrf1^hi272+/−^* and *Tg(fabp10a: CAAX-EGFP)* in *uhrf1^hi272+/−^*background was used for all experiments for ease of liver detection. The live embryos were mounted in 3% methylcellulose on the right side and the left liver lobe was imaged using Nikon Elements on a Nikon SMZ25 Stereoscope. Liver size measurements were calculated in ImageJ using free hand selection of the EFGP/ cherry labeled area corresponding to the left liver lobe, and analyzed using the shape descriptor sub-function to determine the area of the left liver lobe, expressed in µM^2^. For Palbociclib treatment, *uhrf1^−/−^* liver size of 20,000 µM^2^ and below was considered small liver phenotype and *uhrf1^−/−^* liver size above 20,000 µM^2^ considered as rescued.

### Immunofluorescence

120 hpf control siblings and *uhrf1^−/−^* zebrafish embryos containing the *Tg(fabp10a: nls-mcherry)* transgene to identify hepatocyte nuclei were fixed in 4% paraformaldehyde for 4 h at room temperature, washed in PBS, and treated with 150 mM Tris-HCl at pH 9.0 for 5 minutes, followed by heating at 70 °C for 15 minutes according to an established protocol ^93^ and following incubation at room temperature to cool the embryos and washed in PBS. Livers were dissected as described, permeabilized with 10 µg/mL Proteinase K (Macherey-Nagel) in PBS containing 0.1% tween (PBST) for 10 minutes, washed 3 times with PBS, and incubated in a blocking solution containing 5% Fetal Bovine Serum (GIBCO) in PBS for 60 minutes at room temperature. The blocking solution was removed, and the livers were then incubated in 100 μL of H3k9me3 (39161, Activ Motif), H3k9me2 (39753, Activ Motif), Heterochromatin protein (Hp1) (MAB3866, Millipore), or Dnmt1 (SantaCruz) antibody in blocking solution (1:200 dilution) overnight at 4 °C. After 3 washes in PBST, the livers were incubated in secondary antibody (Molecular Probes) in blocking solution (1:400 dilution) in the dark for 2 h on a shaker. After 5 serial washes with PBST, the nuclei were counterstained with Hoechst (Thermo Fisher Scientific) diluted 1:1000 in PBS, washed twice with PBS to remove excess Hoechst, and mounted on a microscope slide with Vectashield (Vector Laboratories) and covered with a 0.1 mM coverslip for imaging using Leica SP8 confocal microscope.

### Terminal Deoxynucleotidyl Transferase dUTP Nick end Labeling Assay (TUNEL)

Larvae collected at 5 dpf were fixed in 4% Paraformaldehyde for 4 h at room temperature, and gradually dehydrated through a graded series of methanol and stored in 100% methanol at 4 ◦C overnight. Gradual rehydration to PBS through a graded series of methanol/PBS dilutions was carried out at room temperature. Larvae were permeabilized with 10 μg/ml Proteinase K (Macherey-Nagel) in PBS containing 0.1% tween (PBST) and fixed in 4% Paraformaldehyde for 10 minutes at room temperature. Livers were then dissected out of the larvae and subjected to TUNEL assay according to manufacturer’s instructions (*In Situ* Cell Death Detection kit, Fluorescein; Roche). Nuclei were counterstained with Hoechst (Thermo Fisher Scientific) diluted 1:1000 in PBS, mounted on a microscope slide with Vectashield (Vector Laboratories) and covered with a 0.1 mM coverslip for imaging using Leica SP8 confocal microscope.

### EdU Incorporation for S-phase Analysis

10 live larvae at 80 hpf, 96 hpf or 120 hpf were transferred into 50 ml falcon tube with 250 uL solution 1 mM EdU (ThermoFisher) in 10% DMSO in embryo water, and incubated on ice for 20 minutes. The tubes were moved to room temperature, filled with embryo water to 50 ml, and incubated for either 12 minutes or 30 minutes at 28.5 °C. After removing the embryo water, the embryos were fixed in 4% Paraformaldehyde (Electron Microscopy Sciences) overnight at 4 °C, then dehydrated through a graded series of methanol and incubated in 100% methanol overnight at 4°C. The embryos were then gradually rehydrated back to 100% PBS by serial dilutions of methanol. The embryos were then permeabilized with PBS containing 1% Triton-X for 15 minutes prior to the Click-iT reaction to attach the Alexa azide (Invitrogen). The larvae were transferred to a solution of 172 µl PBS, 8 µl CuSO4 (100 mM), and 0.2 µl Alexa azide in a 0.6 µl microcentrifuge tube, and rocked in the dark for 10 minutes. 20 µl of ascorbic acid (0.5 M) was added after, and was rocked in the dark for additional 20 minutes. Then the mixture was removed and a fresh Alexa azide solution was added, and the same steps were repeated once more. 80 hpf embryos were individually genotyped by collecting tail samples. The larvae were washed 4 times in PBST prior to dissection and mounted in Vectashield mounting medium with DAPI before they were imaged with the Leica SP8 confocal microscope.

### Confocal Imaging, Image Processing, and Analysis

Confocal imaging was performed using Leica TCS SP8 microscope with a 40x oil immersion at a scan speed of 100 Hz. For antibody immunofluorescence stains and TUNEL experiments, images were acquired at different focal distances from each liver and LAS X software (Leica software) was used for quantification from 3 separate optical sections per liver, which were then averaged from 3 livers per clutch per condition, and a minimum of 2 clutches were analyzed. For EdU and nuclear morphology measurements, Z-stacks were acquired using the galvo stage, with 2 µm intervals. Bit depth was 12, and to enhance image quality, field of view and laser intensity were adjusted separately for each sample. The acquired images were visualized using LASX software. 3D analysis of Z-stacks of Hoechst-stained nuclei was performed using the interactive 3D measurement tool in LASX for volume, surface area, sphericity, elliptical mean, and nuclei counting with Hoechst and EdU stains. Z-stacks were compiled into a 3D image, were adjusted for threshold and noise to define the nuclei clearly, set for minimum size to be measured as 1000 voxels, and measurements of the previously mentioned parameters obtained and exported in excel spreadsheets. For each time point and parameter measured, around 100 nuclei were sampled from at least 3 livers per condition. Results were plotted in GraphPad Prism 9.

### RNA-Seq

50 livers were dissected at 120 hpf from Palbociclib or DMSO treated control sibling or *uhrf1^-/-^*; *Tg(fabp10a: CAAX-EGFP)* larvae as previously described. The livers were individually collected and stored at –80 °C while the carcasses were genotyped for *uhrf1* mutation ^23^. Once genotyped, the *uhrf1^-/-^* livers and control livers from each individual clutch were pooled and RNA extracted according to established protocols ^42^. 100 ng of RNA was used for library preparation according to the manufacturer’s instructions (Illumina, San Diego, CA, USA). Libraries were sequenced on NovaSeq (Illumina) to obtain 150 bp paired-end reads. Raw Fastq files quality was assessed by using FASTQC and the reads were quality trimmed using Trimmomatic ^94^ to remove low quality reads and adapters. Qualified reads were mapped to the reference genome Danio rerio GRCz10. To estimate and compare gene expression in different data sets, raw reads were uploaded to http://tsar.abudhabi.nyu.edu/ for pre-processing and DeSeq2 for Differential gene expression analysis. Datasets of RNAseq performed on 120 hpf *uhrf1^-/-^* and *dnmt1^-/-^*and their controls is previously published ^42^ and available at GEO (GSE160728). Dataset of RNAseq performed on 120 hpf *uhrf1^-/-^* and sibling controls treated with Palbociclib or DMSO will be available on GEO.

### Bioinformatic Analysis

For Gene Ontology and Ingenuity Pathways Analysis (IPA) zebrafish gene names were converted into human gene names using Biomart. Differentially expressed genes were determined by p-value adjusted 0.05 and log 2 fold change 0. GO enrichment analysis of differential expressed genes was conducted using REVIGO ^95^ to find enriched terms for different DEG data sets. An adjusted p-value < 0.05 was considered significant for all analyses. UPSET plots were generated to show intersections of enriched terms from REVIGO, their size, and unique terms ^96^. Heatmaps were generated with Pheatmap on z-score normalized counts. For plotting and statistical analysis, R package ‘ggplot’ and GraphPad Prism 9 software were used. The R codes used are available at https://github.com/SadlerEdepli-NYUAD.

## Statistical Analysis

All experiments were carried out on multiple larvae from at least 2 clutches, as indicated in each figure legend and data represented as the median with range. The unpaired student *t* test was employed to compare the medians between groups with adjustment for multiple comparisons and all of the data were considered to be significant at \**p* < 0.05, \*\**p* < 0.005, \*\*\**p* < 0.0005. All plots along with statistical analysis were generated in GraphPad Prism 9 (GraphPad Software). Bioinformatic analysis and visualization of genomic data were performed and plotted in RStudio.

## Author Contributions

Conceptualization, K.C.S., B.M.; methodology, B.M., E.M., S.R.; formal analysis, B.M., E.M., S.R.; investigation, K.C.S., B.M., E.M., S.R.; writing—original draft preparation, K.C.S. and B.M., writing—review and editing, K.C.S., B.M., E.M.; visualization, K.C.S., B.M., E.M., S.R.; supervision, K.C.S.; project administration, K.C.S.; funding acquisition, K.C.S. All authors have read and agreed to the published version of the manuscript.

## Data Availability Statement

All the datasets used in this paper are available as GEO datasets under the number GSE160728. Code used in this manuscript is available on https://github.com/SadlerEdepli-NYUAD.

## Acknowledgments

FACS Cell sorting was performed in the NYUAD CGSB FACS Core at New York University Abu Dhabi. Bioinformatics assistance was provided by the NYUAD Bioinformatics Core. We are grateful to Mehar Sultana, Marc Arnoux, Nizar Drou and Muhammad Arshad for assistance with sequencing and analysis Mehar Sultana for assistance with technical support for the FACS experiments, and Rachid Rezgui for assistance with NYUAD Light Microscopy Core Technology Platform. All members of the Sadler group provided insightful discussion and help throughout the project, in particular Patrice Delaney bioinformatics and editing assistance.

## Supplemental Materials

### Supplementary materials and methods

#### Survival assay

Heterozygous *uhrf1^hi272+/-^*; *Tg(fabp10a: nls-mcherry) ^+/-^* adults were incrossed and split into 40 embryos per 90 mm petriplate, with around 25% *uhrf1^-/-^* per plate. The embryos were dechorionated and treated with either 0.5% DMSO or 20 μm Palbociclib at 48 hpf. At 120 hpf, the embryos were transferred to fresh embryo water and monitored till 9 dpf when the live embryos were individually collected and genotyped for the *uhrf1* mutation.

#### Eye size measurement

Eye size measurements were carried out based on the 2-dimensional area of the left eye in 120 hpf larvae. Live larvae were mounted in 3% methylcellulose on the right side and the left eye was imaged using Nikon Elements on a Nikon SMZ25 Stereoscope. Eye size measurements were calculated in ImageJ using free hand selection of the eye area and analyzed using the shape descriptor sub-function to determine the area expressed in µM^2^.

## Supplement Figure Legends

**Figure S1.** Representative images of 5 minute, 12 minute and 30 minute EdU incorporation in 120 hpf *uhrf1* control and *uhrf1^-/-^*livers.

**Figure S2.** Phenotype of Palbociclib (PD) treated *uhrf1* control and mutants (A) 120 hpf embryos from incross of *uhrf1^+/-^; Tg(fabp10a: nls-mcherry)* adults treated with DMSO and imaged, livers are in red. (B) Phenotypic assessment of the embryos at 120 hpf. (C) 120 hpf embryos from incross of *uhrf1^+/-^; Tg(fabp10a: nls-mcherry)* adults treated with PD and imaged, livers in red, embryos numbered and genotype on the right side of the image. (D) PCR based genotyping to ascertain the *uhrf1^-/-^* with PD treatment. (E) Eye size measurement of 120 hpf *uhrf1^-/-^* and control embryos with DMSO and PD treatment. (F) Survival of DMSO and PD treated embryos at 9 dpf. Scale: 50µm, the number of samples and clutches indicated for each condition. p-value *< 0.05, **< 0.005, *** < 0.0005 by two tailed Students t-test with adjustment for multiple comparisons.

**Figure S3.** Palbociclib (PD) treatment differentially regulates cell cycle gene expression but does not rescue transposon expression in *uhrf1* mutants (A) Volcano plot of Differently Expressed Genes (DEGs) from RNA-Seq between 120 hpf *uhrf1* control PD treated and control DMSO treated livers. (B) IPA showing the pathways differentially expressed from RNA-Seq between 120 hpf *uhrf1* control PD treated and control DMSO treated livers. (C) Four quadrant cross-plot of the DEGs with log2fold change from RNA-Seq between 120 hpf *uhrf1^-/-^* PD treated and DMSO treated livers. (D) Correlation plot of repetitive elements in 120 hpf *uhrf1*^−/−^ mutant livers treated with DMSO and PD. Upregulated TEs have log 2 fold change > 0; downregulated TEs have log 2 fold change < 0. Log 2 fold change is calculated between mutants and their own control siblings. (E) Gene ontology (REVIGO) of the upregulated and downregulated genes in 120 hpf DMSO and PD treated *uhrf1^-/-^* liver RNA-Seq using significant genes (*p_adj_*< 0.05) for each category.

**Figure S4.** Palbociclib (PD) treatment on *uhrf1* mutants rescues key cell cycle gene overexpression (A) IPA analysis of the G1-S pathway in DMSO *uhrf1^-/-^* compared to DMSO treated controls. The color of the circles represents the observed results (red for activated/ upregulated and blue for repressed/ downregulated). (B) qPCR of key cell cycle genes of 120 hpf *uhrf1^-/-^*and control livers treated with DMSO or PD, data from 2 clutches shown. (C) Volcano plot of Differently Expressed Genes (DEGs) from RNA-Seq between 120 hpf *uhrf1^-/-^*PD treated and *uhrf1^-/-^* DMSO treated livers. (D) IPA analysis of the G1-S pathway between PD and DMSO treated *uhrf1^-/-^*. The color of the circles represents the observed result (red for activated/ upregulated and blue for repressed/ downregulated).

**Figure S5.** IPA analysis of ATM pathway IPA analysis of the ATM pathway in 120 *uhrf1^-/-^* liver and 120 *dnmt1^-/-^* liver transcriptomes. The color of the circles represents the observed results (orange for activated/ upregulated only in *uhrf1^-/-^*, red for activated in both, light blue for repressed/ downregulated only in *uhrf1^-/-^* and dark blue for repressed in both.

**Figure S6.** Testing the efficiency of *dnmt1^cr^ in c269off+/-; 14X UAS:dsRed+/-; Tg(fabp10a:Gal4;cmlc2:GFP)+/-* and nuclear morphology of *dnmt1^c^*^r^ injected *uhrf1* control and mutants (A) Representative images of 120 hpf uninjected and *dnmt1^cr^* embryos imaged for dsRed and GFP in the head. GFP positivity indicates hypomethylation. (B) 120 hpf embryos from incross of *uhrf1^+/-^*;*Tg(fabp10a: nls-mcherry)* adults injected or uninjected with *dnmt1^cr^* were genotyped, PFA fixed, livers dissected, Hoechst stained and imaged as a Z-stack which were then used to calculate (B) volume, (C) Surface area and (D) Ellipsoid mean of the hepatocytes. The number of samples and clutch indicated for each condition. p-value *< 0.05, **< 0.005, *** < 0.0005 by ANOVA.

## Supplemental tables

Table S1. REVIGO analysis of RNA-Seq from 120 hpf *uhrf1* mutant livers

Table S2. Cell cycle pathways identified by IPA of RNA-Seq from 120 hpf *uhrf1* mutant livers

Table S3. Optimization of PD exposure protocol on WT embryos

Table S4. DEGs in the liver of PD and DMSO treatment of 120 hpf control larvae

Table S5. IPA of DEGs from the liver of PD and DMSO treated 120 hpf control larvae

Table S6. DEGs from the liver of DMSO treated control and *uhrf1* mutant larvae and from PD treated control and *uhrf1* mutants at 120 hpf

Table S7. ENSEMBL IDs of the heatmap clusters from DMSO and PD treated control and *uhrf1* mutant larvae

Table S8. REVIGO analysis of clusters 4 and 5 from DMSO and PD treated control and *uhrf1* mutant larvae

Table S9. REVIGO analysis of DEGs between *uhrf1* mutants treated with DMSO or PD (from Table S6)

Table S10. DNA replication DEGs, immune DEGs and TNFa DEGs between *uhrf1* mutants treated with DMSO or PD

Table S11. DEGs from the liver of DMSO vs. PD treated *uhrf1* mutants at 120 hpf

Table S12. IPA of DEGs from DMSO vs. PD treated *uhrf1* mutants at 120 hpf

Table S13. Primers for qPCR and CRISPR sgRNA

