## Supplementary figures and images for "DNA hypomethylation activates Cdk4/6 and ATR to cause dormant origin firing and cell cycle arrest that restricts liver outgrowth in zebrafish"

### Figure S1

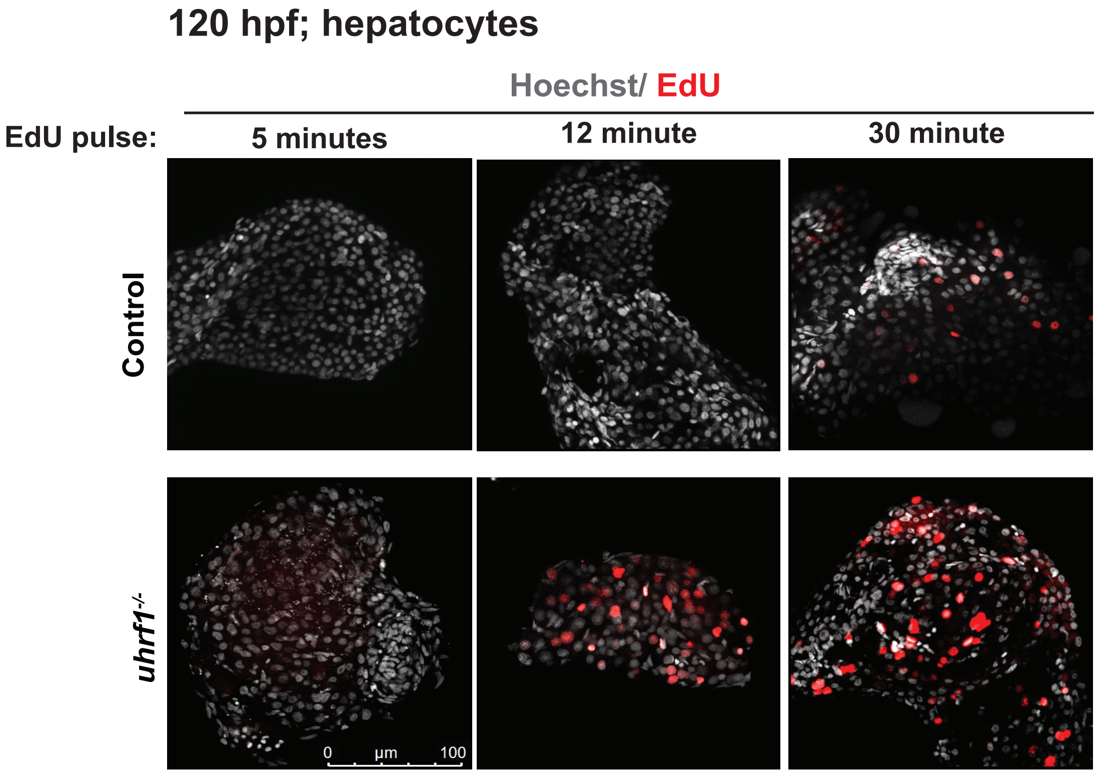

### Figure S2

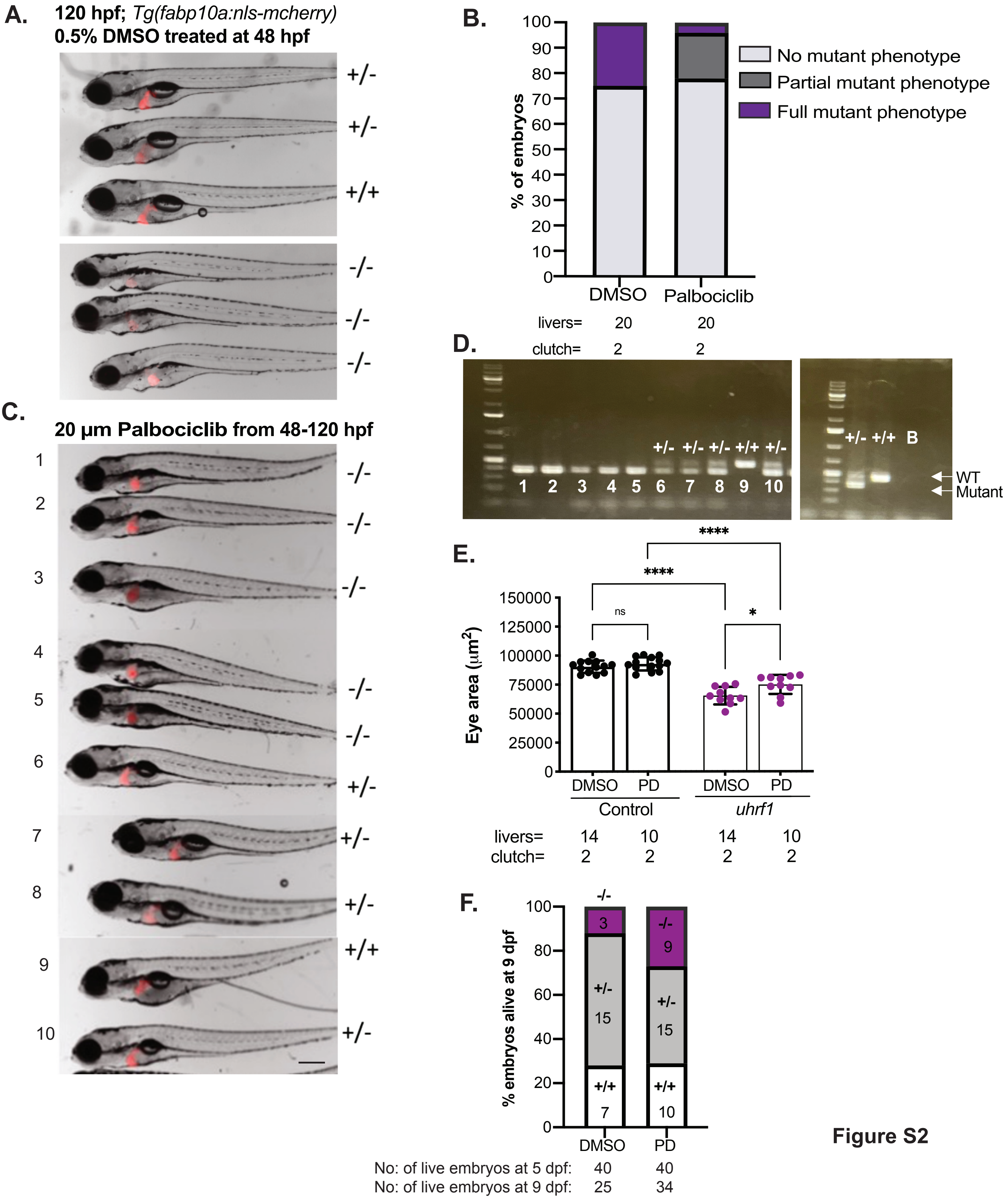

### Figure S3

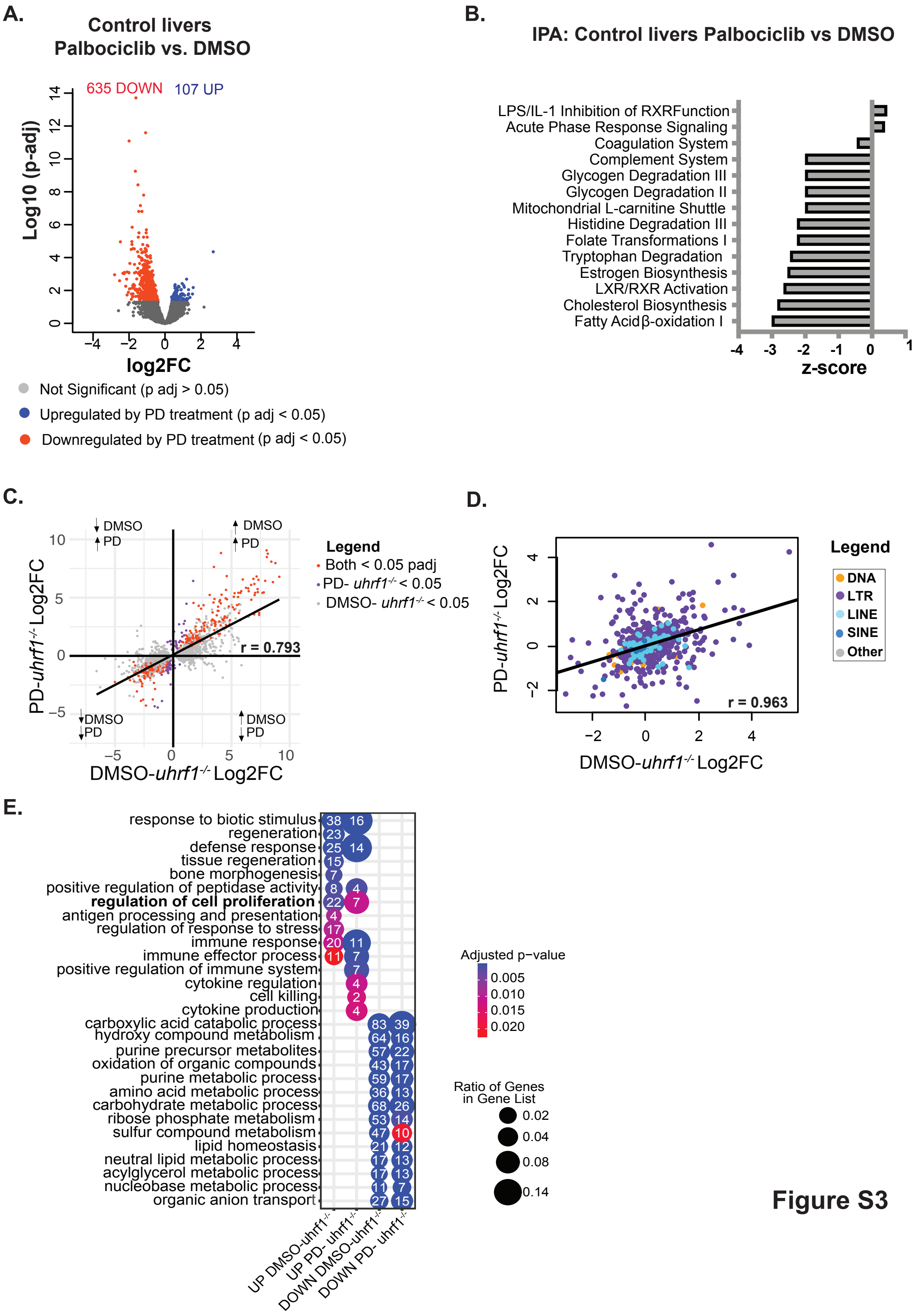

### Figure S4

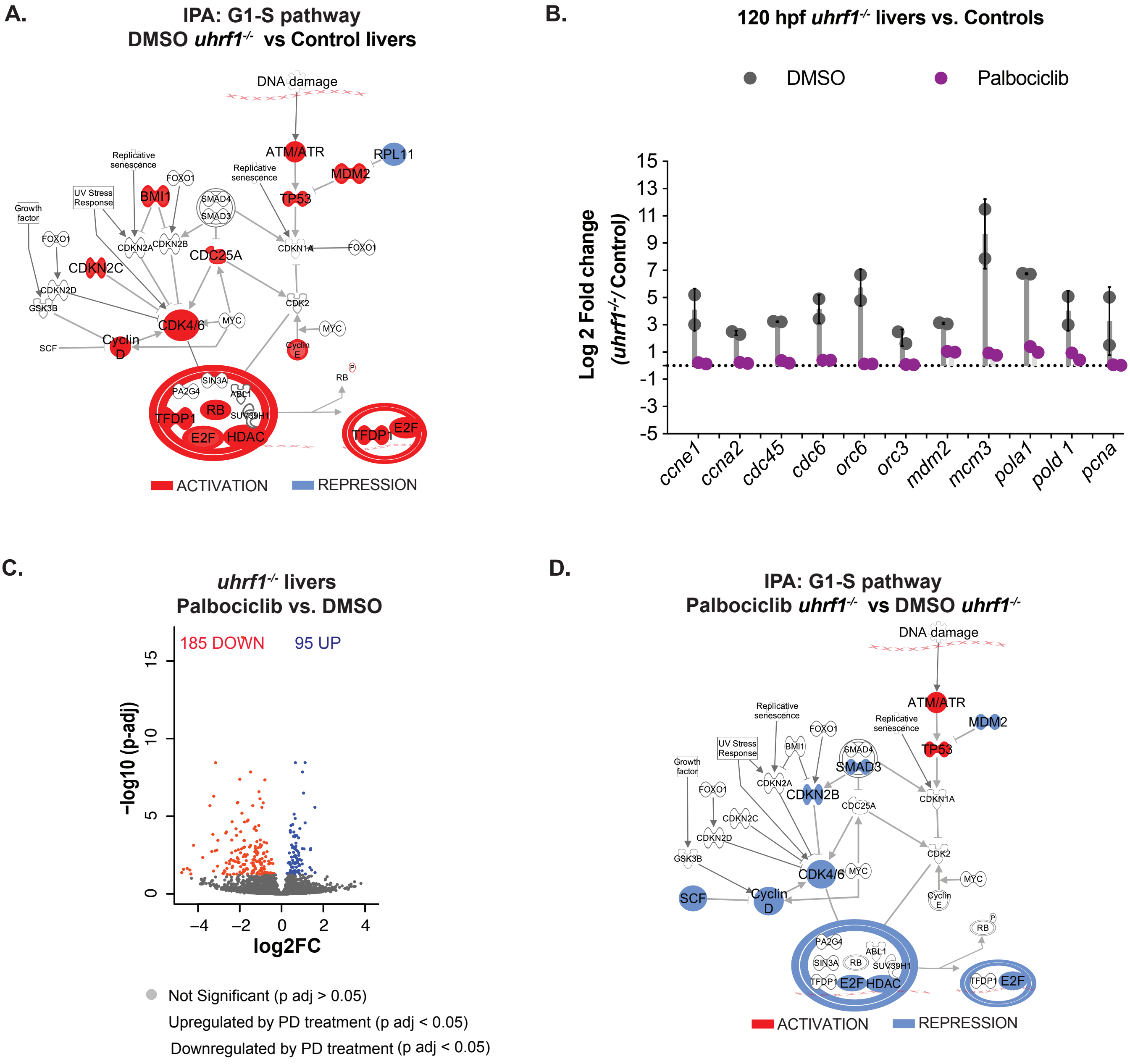

### Figure S5

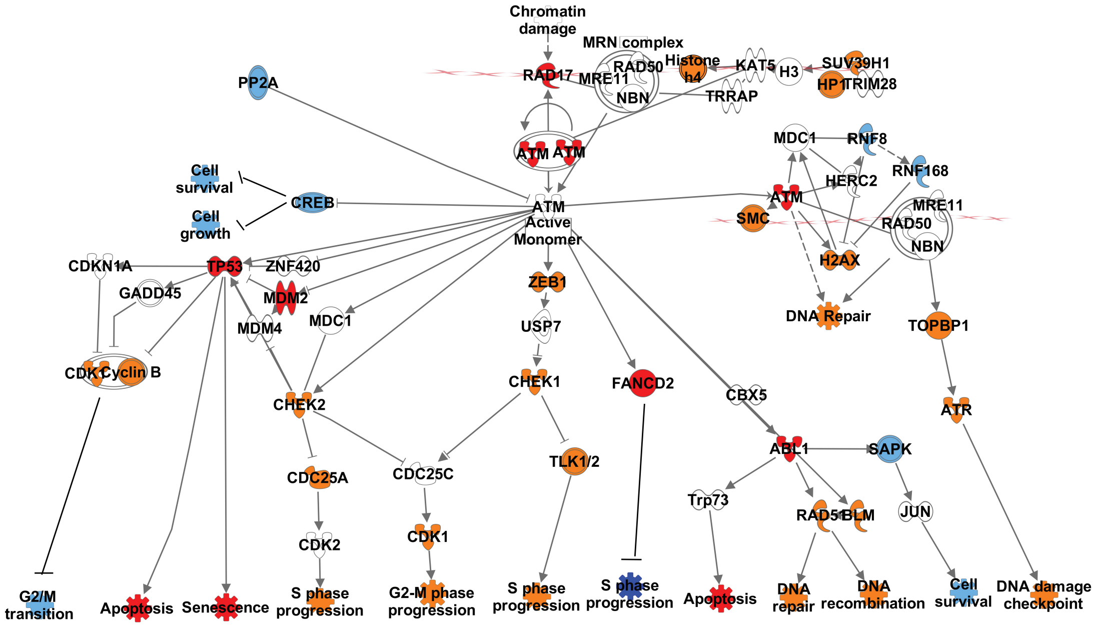


#### Legend

- UP *uhrf1*$^{-/-}$ only
- DOWN *uhrf1*$^{-/-}$ only
- UP *uhrf1*$^{-/-}$ and *dnmt1*$^{-/-}$
- DOWN *uhrf1*$^{-/-}$ and *dnmt1*$^{-/-}$

**Figure S5**

### Figure S6

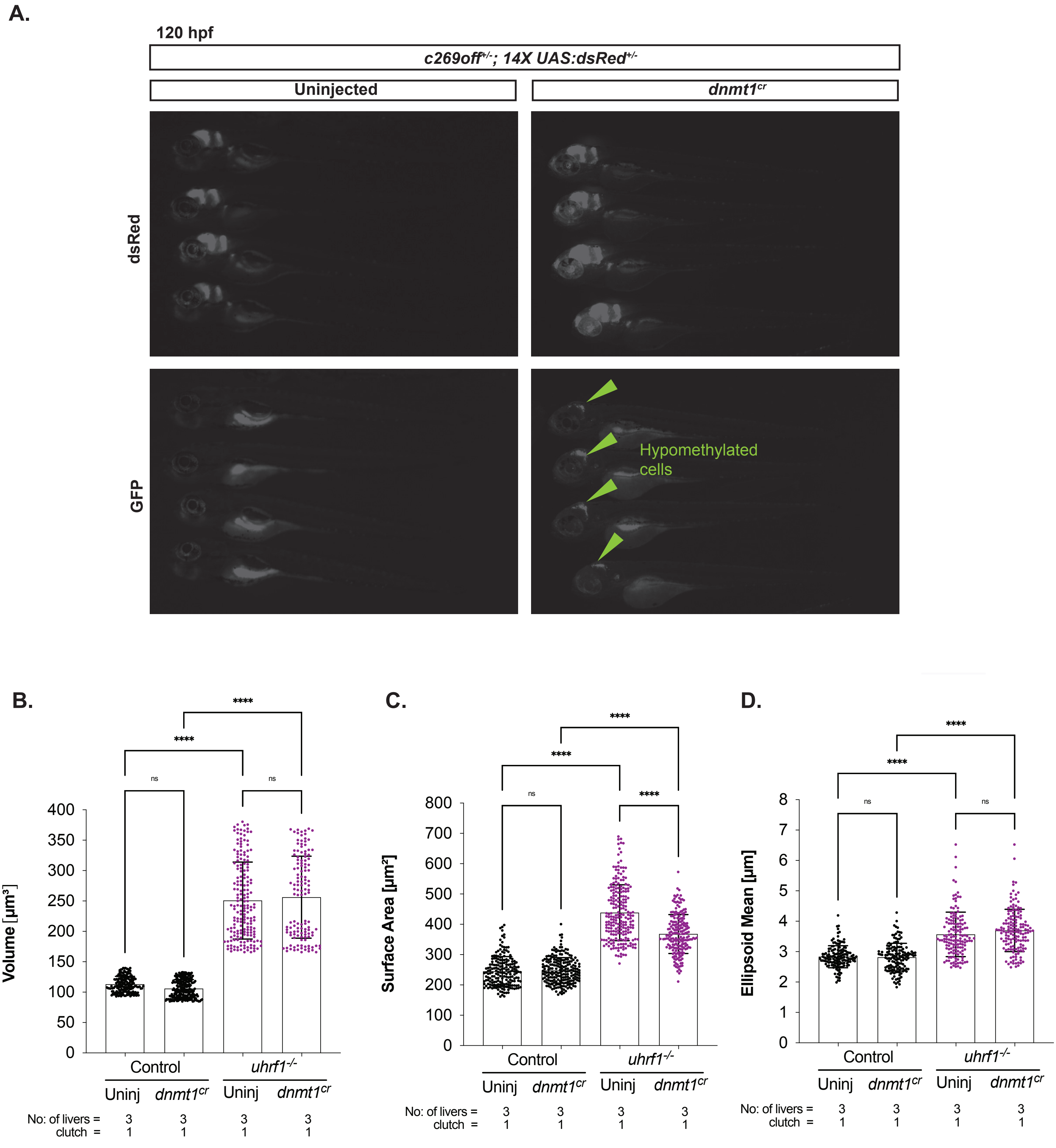
